# Svep1 stabilizes developmental vascular anastomosis in reduced flow conditions

**DOI:** 10.1101/2021.03.13.435246

**Authors:** Baptiste Coxam, Yvonne Padberg, Katja Maier, Simone Jung, Eireen Bartels-Klein, Anna Szymborska, Lise Finotto, Christian S.M. Helker, Didier Y.R. Stainier, Stefan Schulte-Merker, Holger Gerhardt

## Abstract

We report the discovery that flow and Svep1 are modulator of vessel anastomosis during developmental angiogenesis in zebrafish embryos. We show that loss of Svep1 and blood flow reduction both contribute to defective anastomosis of intersegmental vessels. We show that this defect in primary angiogenic sprouts is associated with an expansion of Apelin-positive tip cells and with reduced formation and lumenisation of the dorsal longitudinal anastomotic vessel. Mechanistically, our results suggest that flow and Svep1 act synergistically to modulate vascular network formation in the zebrafish trunk.

## Introduction

Angiogenesis defines the formation of new vessels from pre-existing ones and is a stepwise process that leads to the formation of a perfused network of arteries and veins that are optimally organised to serve the metabolic needs of the developing embryo. In zebrafish embryos, vasculogenesis results in the formation of the two main blood vessels, namely the dorsal aorta (DA) and the posterior cardinal vein (PCV). Later, angiogenic sprouts emerge from the DA. These multicellular sprouts are composed of a leading tip cell followed by stalk cells [1], and migrate dorsally in between the vertical somite boundaries to form intersegmental vessels (ISVs)[2]. The proliferation and migration of ISVs to the dorsal part of the embryo is driven by Vegfa signalling, through binding to the VEGFR2 zebrafish ortholog Kdr and its ohnolog Kdrl [3–8]. Activation of Kdrl signalling in the leading ISV endothelial cell (tip), mediated in large part via downstream phosphorylation of the serine/threonine kinase ERK1/2, promotes migratory behaviour. In parallel, activation of Kdrl signalling in the tip cell leads to Notch mediated inhibition of Kdrl signalling in trailing cells (stalks), preventing their conversion into tip cells and promoting their proliferation to support ISV expansion [6, 9, 10]. Around 30-32 hours post-fertilisation (hpf), the leading tip cells of the ISVs start to anastomose with their ipsilateral neighbours, a process that ultimately leads to the formation of the dorsal longitudinal anastomotic vessel (DLAV), dorsal to the neural tube [11, 12]. The DLAV is initially a paired bilateral structure that is fully lumenised by 48 hpf, but subsequently both sides progressively connect to form a complex plexus [2, 12]. Zygmunt and colleagues demonstrated that maturation of the DLAV plexus is regulated by flow and Vegfr signalling after 48 hpf. However, while they show that flow is dispensable for the initial formation of the DLAV, little is known about the cellular mechanism driving the anastomosis of ipsilateral ISVs and the lumenisation of the DLAV segments during DLAV formation (32 to 48 hpf).

Recent reports have characterised the importance of *svep1/polydom*, a secreted ECM protein that has been reported to mediate cell-to-substrate adhesion in vitro, in an integrin α9β1-dependent manner [13], as a regulator of secondary angiogenesis [14, 15]. In zebrafish, loss of function *svep1* mutants exhibit a reduced number of venous and lymphatic precursor (parachordal lymphangioblasts, PLs) emerging from the PCV during secondary angiogenesis (from 32 hpf). In addition, PLs show reduced migration capacity from the horizontal myoseptum. Both these defects lead to an increased number of arterial ISVs (aISVs) and a severe reduction in lymphatic trunk vasculature. Here, we uncover a completely new and distinct role for Svep1 in the regulation of DLAV formation under reduced flow conditions, acting in part through modulation of Vegfa/Vegfr signalling in endothelial cells.

## Material and Methods

### Zebrafish husbandry and transgenic lines

Zebrafish (*Danio rerio*) were raised and staged as previously described [16]. The following transgenic lines were used: *Tg[fli1a:EGFP]^y1^* [17] (labeling all endothelial cells), *Tg[gata1a:dsRed]^sd2^* [18] (labeling all erythrocytes), *Tg[-0.8flt1:RFP]^hu5333^* [5] (strongly labels arterial endothelial cells)*, TgBAC(apln:eGFP)^bns157^* [19] *(labeling endothelial tip cells)*. Tg(svep1:Gal4FF;UAS:GFP) (labeling *svep1* positive cells) [14]. The *svep1^hu512+/-^* line has been described previously [14]. For growing and breeding of transgenic lines we comply with regulations of the ethical commission animal science of MDC Berlin and with FELASA guidelines [20].

### Tricaine treatment

To slow down heart rate and blood flow during DLAV formation, embryos were treated with 0.007%, 0.014% (1X) or 0.028% (2X) tricaine (MS-222, Sigma) between 30 and 48 hpf, as indicated in the figure legends.

### Morpholino knockdown

Morpholinos against *svep1* (5ng) and *flt1* (1ng) were used as previously described in [15, 21] and injected in the yolk of zebrafish embryos at the one-cell stage.

### Statistical Analysis

All quantifications were performed in the trunk region of zebrafish embryos, across 7-9 somites. (N) refers to experimental replicates, (n) refers to number of embryos. Statistical analysis was performed with Mann-Whitney-U tests, unless indicated otherwise. No statistical method was used to predetermine sample size. Data represent mean ± standard deviation of representative experiments (except when indicated otherwise). Statistical tests were conducted using Prism (GraphPad) software. Adequate tests were chosen according to the data to fulfil test assumptions. Sample sizes, number of repeat experiments, performed tests and p-values are indicated per experiment.

Zebrafish embryos were selected based on the following pre-established criteria: normal morphology, beating heart, presence of circulating red blood cells. The experiments were not randomized. For every experiment treated and control embryos were derived from the same egg lay. The investigators were not blinded to allocation during experiments and outcome assessment.

### Live imaging

Embryos were anaesthetized in 0.014% tricaine (MS-222, Sigma), mounted in a 35 mm glass bottom petri dish (0.17 mm, MatTek) using 0.6-1% reduced melting point agarose (Sigma) containing 0.014% tricaine, and bathed in E3 media containing 0.014% (1X tricaine) and 0.003% PTU. Time-lapse imaging was performed using an upright 3i spinning-disc confocal using a Zeiss Plan-Apochromat, 20x, 40x/1.0 NA water-dipping objective. Image processing was performed using Fiji software [22].

### Isolation of endothelial cells

48 hpf *Tg[fli1a: nEGFP]^y7^* and *Tg[fli1a: EGFP]^y1^* crossed to *Tg[gata1:dsRed]* embryos were dechorionated using a solution of 1mg/mL Pronase (Sigma Aldrich) on an orbital shaker for 10 minutes at room temperatures. Up to 250 dechorionated embryos per conditions were anaesthetized with 1X (0.014%) tricaine and transferred to a 1.5 mL Eppendorf with 1 mL calcium-free Ringer solution (116 mM NaCL, 2.9 mM KCL, 5 mM HEPES pH7.2) to remove the yolks. After pipetting gently up and down with a 1mL tip the embryos were centrifugated at 2000 rpm for 5 minutes at 4°C. The supernatant was removed, and the procedure repeated until all the yolks were removed and the solution clear. The calcium-free ringer solution was replaced with 1 mL of protease solution (72 μg/mL Liberase DH research grade from Merck/Sigma, 0.4U/mL DNaseI-Invitrogen). The embryos were incubated at 28.5C on an orbital shaker for 20 minutes, pipetting up and down with a 200μL tip every 3 minutes, to form a homogenous solution of cells. The dissociation process was stopped by placing the embryos on ice and adding 2μL CaCl2 and 0.5 μL FBS per mL. The cell suspension was centrifuged for at 2000 rpm for 5 minutes at 4°C. The supernatant was discarded, and the cells resuspended in sorting solution (2mM EDTA, 0.4U/mL DNaseI, 0.5% FBS in DPBS). The solution was passed through a 40μm strainer inside a 50 mL falcon tube, previously washed with 500μL sorting solution. Following filtration, 500μL sorting solution was added to the strainer. The filtered solution was centrifuged at 2000 RPM for 5 minutes at 4C. The supernatant was removed, and the cell resuspended in 700 μL sorting solution. The cell suspension was then loaded onto an ARIA III FACsorter (BD Bioscience). Using *Tg[gata1:dsRED]* only embryos for gating, we specifically sorted GFP+, dsRED-cells to remove red blood cells with *fli1a* promoter activation at this specific developmental stage. Upon centrifugation and removal of the supernatant, all cells were stored immediately at -80C until protein extraction.

### Protein extraction

Sorted embryos were treated with 40μL Lysis Buffer (1mL 1M Tris-HCl, 0.4 mL 0.5M EDTA, 8.75 mL 10% Brij 96, 1.25 mL 10% NP-40 to 100mL with dH2O) and 0.4μL Protease Inhibitor cocktail (ThermoFischer). The samples were homogenized with a pestle and centrifuged at 13000 RPM for 15 minutes at 4C. The protein supernatant was collected, and protein concentration assessed using a BCA protein assay kit (ThermoFischer).

### Western Blot

20-50μg of the protein lysates (equal amount for each condition to compare) were diluted in 18.7μL water and 6.3μL loading buffer (25μL total volume) and heated for 5 min at 95°C to denature proteins. The samples were loaded and run alongside a 10μl ladder marker (Novex Sharp Pre-Stained -thermoFischer) for 1h at 150V and subsequently transferred onto previously MeOH activated polyvinylidene fluoride membranes. Membranes were blocked with 5% non-fat dry milk in 50 mg/mL TBS-T for 1.5h at room temperature and then incubated with primary antibody against p-ERK, overnight at 4°C (1:250, *(Erk1/2) (Thr202/Tyr204)* #9101, Cell Signaling Technology).

After incubation with primary antibody, the membranes were washed 4 times in 50 mg/mL TBS-T and then incubated with secondary antibodies (1:4000 anti-rabbit) and washed 3 times with 50 mg/mLTBS-T. Immunodetection was performed using a chemiluminescence kit (1mL SuperSignal West Dura; Pierce), and bands were developed using the Las-4000 imaging system. After initial immunodetection, membranes were stripped of antibodies by using the Stripping kit (ThermoFisher) at 56°C for 40min and re-probed with anti–GFP antibody for 1h (1:1000, Origene R1091P). Band intensity was measured using the histogram function on the Fiji software, with control and treated samples on the same blot [22].

### Blood flow and heart rate measurements

Embryos were anaesthetized and imaged on an upright 3i spinning-disc confocal using a Zeiss Plan-Apochromat, 20x/1.0 NA water-dipping objective with a frame interval of 10ms. Kymographs were generated using the MultipleKymograph plugin in ImageJ to quantify heart rate over an 8 second period, synced to the beginning of a heartbeat (line width: 1).

To estimate instantaneous blood flow speed, we cropped images of the dorsal aorta and measured average frame-to-frame translation of red blood cells using the Kuglin-Hines algorithm (Kuglin and Hines, 1975) for image phase-correlation. In brief, the phase correlation map between two adjacent frames was calculated by multiplying the Fast Fourier transform (FFt) of frame_i_ and a conjugate FFt of frame_i+1_. The inverse FFt of the phase correlation provides a correlation map with a peak offset from the center by the relative shift between the frames. The position of the peak was determined by finding the local maximum in a Gaussian filtered correlation map. The velocity data was smoothed with a moving average filter with a span of 5 frames. Analysis was performed in Matlab (Mathworks, Inc.).

### Chemical treatment

Where indicated, embryos were treated with VEGFR inhibitors ZM323881 (Tocris Bioscience) or SU5416 (sigma Aldrich), from 30 to 48hpf, in addition to 0.003% PTU and the indicated amount of tricaine.

## Results

### *svep1* mutant and morphant zebrafish embryos exhibit vascular anastomosis defects

Imaging angiogenesis in the trunk of *svep1* loss-of-function mutants (Ly02-12 – named *svep1^hu4767^* in this manuscript) [14] from 30 to 48 hpf, we noticed that a significant number of primary angiogenic sprouts failed to anastomose with their ipsilateral neighbours (Figure 1A) (Supplementary Video 1, 2). In the majority of cases (74% ± 33), the DLAV gaps arise following the regression of a pre-existing connection between ipsilateral neighbouring ISVs, rather than an absence of connection (N=6 experiments, n=18 mutants). Additionally, only a minority of these connections (13%, N=6 experiments, n=17 mutants) were transiently lumenised before regressing. These results suggest that in this context, *svep1* loss of function negatively affects the stabilisation of vascular connections between neighbouring sprouts.

**Figure 1.**
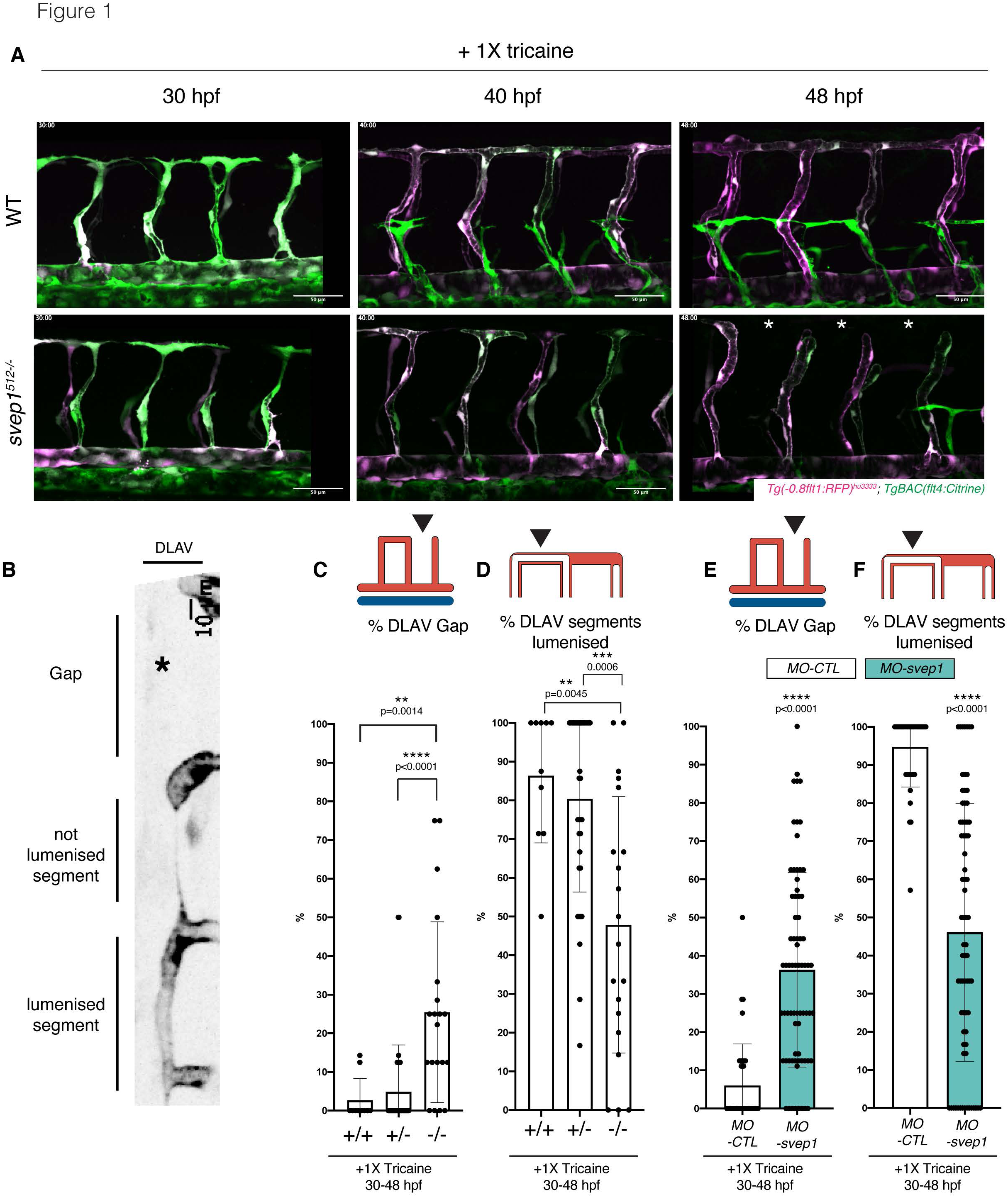
*svep1* mutants and morphants zebrafish embryos exhibit vascular anastomosis defects. **(A)** Stills from time lapse movie of *MO-CTL* (5ng) and *MO-svep1* (5ng) *Tg(−0.8flt1:RFP)^hu3333^; TgBAC(flt4:Citrine)* embryos treated with 1X (0.014%) tricaine from 30 to 48 hpf. White asterisks indicate gaps in the DLAV. **(B)** Still from a time lapse movie of *MO-CTL* (5ng) *Tg(−0.8flt1:RFP)^hu3333^; TgBAC(flt4:Citrine)* embryo exhibiting a gap in the DLAV between two adjacent ISVs. Side view, dorsal side left. **(C)** Bilateral quantifications of the percentage of gaps in the DLAV at 48 hpf in *svep1*^512^ WT (n=5), heterozygous (n=18) and homozygous mutants (n=10) treated with 1X tricaine (0.014%) from 30 to 48 hpf (N=3). **(D)** Bilateral quantifications of the percentage of lumenised segments in the DLAV at 48 hpf in *svep1*^512^ WT (n=5), heterozygous (n=18) and homozygous mutants (n=10) treated with 1X tricaine (0.014%) from 30 to 48 hpf (N=3). (**E**) Bilateral quantifications of the percentage of gaps in the DLAV at 48 hpf in *MO-CTL* (5 ng) (n=20) and *MO-svep1* (5ng) (n=36) embryos treated with 1X tricaine (0.014%) from 30 to 48 hpf (N=6). (**F**) Bilateral quantifications of the percentage of lumenised segments in the DLAV at 48 hpf in *MO-CTL* (5 ng) (n=20) and *MO-svep1* (5ng) (n=36) embryos treated with 1X tricaine (0.014%) from 30 to 48 hpf (N=6).

We quantified the number of gaps in the DLAV, and lumenisation status of existing DLAV segments at 48 hpf, a time at which the DLAV is considered fully formed and almost fully lumenised in wild type (WT) zebrafish embryos [2, 12]. *svep1*^512^ mutants exhibited a significantly increased number of gaps in their DLAV at 48 hpf (25.5% ± 23.4 versus 2.6% ± 5.6 in their WT siblings) and a significant decrease of lumenised DLAV segments (47.8 % ± 33.1 versus 86.4 % ± 17.3) (Figure 1B-D). These phenotypes were also observed in *svep1* morphants compared to control morphants (DLAV gaps: 36.3% ± 25.4 versus 6% ± 10.9; lumenised DLAV segments: 46.1% ± 33.8 versus 94.7 ± 10.5) (Figure 1E, F). In addition, while the expressivity of the morphant phenotype in different zebrafish transgenic lines and clutches varied markedly, we observed a statistically significant difference between *svep1* and control morphants in all cases, (Supplementary Figure 1).

### *svep1* loss-of-function sensitises angiogenic remodelling to reduced blood flow

Surprisingly, we could not detect any anastomosis defects at the DLAV in *svep1^512^* mutants and morphants when imaged at 2 dpf, whilst they continued to exhibit the previously reported PL phenotypes [14] (PLs at the horizontal myoseptum: 47.6% ± 17.5 versus 80.2% ± 19.4% in WT clutch mates and aISVs: 65.4% ± 15.1 versus 49.3% ± 15 in WT siblings, N=4, n=12 mutants, n=25 WT)). The DLAV phenotypes instead only occurred in mutant embryos that were live imaged from 30 to 48 hpf. Upon closer inspection, we found that treatment with tricaine (tricaine mesylate – MS222), a muscle relaxant commonly used to immobilise zebrafish embryos, lead to a dose-dependent emergence of the DLAV phenotypes in *svep1* loss-of-function morphants (Figure 2A, B) following treatment with concentration of 1X (0.014%) or above from 30 to 48 hpf. In addition, removal of tricaine from mutant embryos lead to a significant recovery of the DLAV vasculature at 72 hpf, with only 27% (± 26) of the DLAV gaps still present at that stage (supplementary Figure 2A, B).

**Figure 2.**
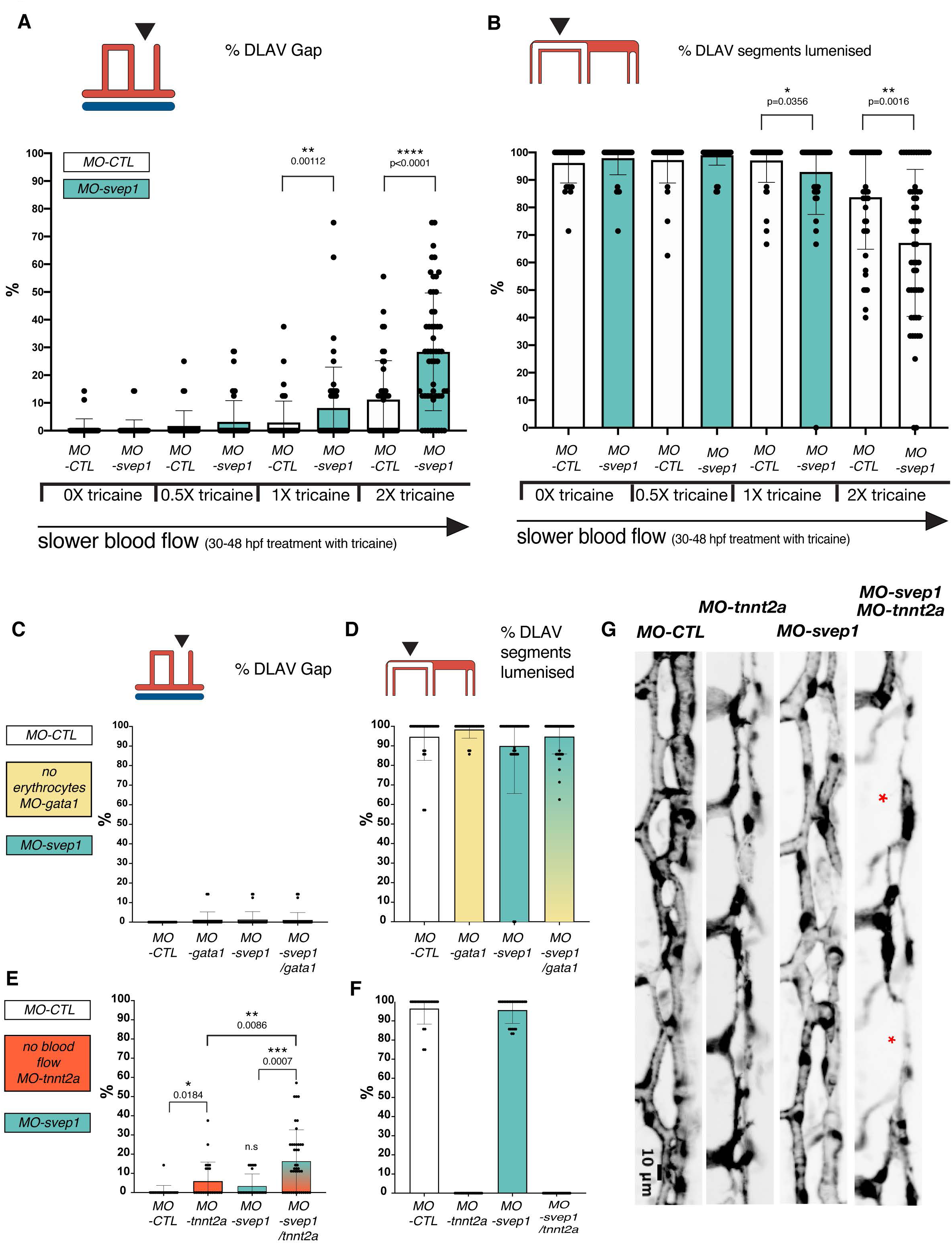
*svep1* loss-of-function sensitises angiogenic remodelling to reduced blood flow. **(A)** Bilateral quantifications of the percentage of gaps in the DLAV at 48 hpf in *MO-CTL* (5 ng) and *MO-svep1* (5ng) embryos treated with 0X (n=14 *MO-CTL*, n=20 *MO-svep1*), 0.5X (0.007%) (n=16 *MO-CTL*, n=24 *MO-svep1*), 1X (0.014%) (n=22 *MO-CTL*, n=27 *MO-svep1*) or 2X (0.028%) (n=21 *MO-CTL*, n=27 *MO-svep1*) tricaine from 30 to 48 hpf (N=3). **(B)** Bilateral quantifications of the percentage of lumenised segments in the DLAV at 48 hpf in *MO-CTL* (5 ng) and *MO-svep1* (5ng) embryos treated with 0X (n=14 *MO-CTL*, n=20 *MO-svep1*), 0.5X (0.007%) (n=16 *MO-CTL*, n=24 *MO-svep1*), 1X (0.014%) (n=22 *MO-CTL*, n=27 *MO-svep1*) or 2X (0.028%) (n=21 *MO-CTL*, n=27 *MO-svep1*) tricaine from 30 to 48 hpf (N=3). **(C)** Bilateral quantifications of the percentage of gaps in the DLAV at 48 hpf in *MO-CTL* (5 ng) (n=13), *MO-gata1* (8ng) (n=12), *MO-svep1* (5ng) (n=16) and *MO-gata1* (8ng)/*MO-svep1* (5ng) (n=25) embryos (N=3). **(D)** Bilateral quantifications of the percentage of lumenised segments in the DLAV at 48 hpf in *MO-CTL* (5 ng) (n=13), *MO-gata1* (8ng) (n=12), *MO-svep1* (5ng) (n=16) and *MO-gata1* (8ng)/*MO-svep1* (5ng) (n=25) embryos (N=3). **(E)** Bilateral quantifications of the percentage of gaps in the DLAV at 48 hpf in *MO-CTL* (5 ng) (n=11), *MO-tnnt2a* (4ng) (n=12), *MO-svep1* (5ng) (n=12) and *MO-tnnt2a* (4ng)/MO-svep1 (5ng) (n=21) embryos (N=3). **(F)** Bilateral quantifications of the percentage of lumenised segments in the DLAV at 48 hpf in *MO-CTL* (5 ng) (n=11), *MO-tnnt2a* (8ng) (n=12), *MO-svep1* (5ng) (n=12) and *MO-tnnt2a* (4ng)/*MO-svep1* (5ng) (n=21) embryos (N=3). **(G)** Maximum intensity projection of representative DLAV in *MO-CTL* (5 ng) (n=11), *MO-tnnt2a* (8ng) (n=12), *MO-svep1* (5ng) (n=12) and *MO-tnnt2a* (4ng)/MO-svep1 (5ng) (n=21) embryos at 48 hpf. Red asterisks indicate gaps.

Previous work revealed that svep1 loss of function is associated with cardiac defects [23]. To test if *svep1* loss-of-function augments tricaine-induced blood flow reduction, we quantified heartbeats per minutes in control and svep1 morphants and found no differences in embryos treated with 1X (0.014%) or 2X (0.028%) tricaine from 30 to 48 hpf (supplementary Figure 2C). However, we found the mean blood flow speed to be significantly decreased in svep1 morphants compared to control morphants when treated with 1X tricaine (MO-CTL 5ng: 679um/s ± 205, MO-svep1 5ng: 521 um/s ± 190). This suggests that the cardiac phenotype reported in *svep1* loss-of-function embryos exacerbates the blood flow reduction induced by tricaine treatment.

To further characterize the phenotype, we therefore decided to begin our investigation at 30 hpf, as this time point marks the beginning of ISV ipsilateral anastomosis in most embryos (Figure 1A). As tricaine treatment leads to a reduction of blood flow speed [24], we investigated whether a general blood flow speed reduction or a reduction of erythrocytes-dependent shear stress is responsible for the DLAV phenotype in *svep1* morphants. In the absence of tricaine treatment, simultaneous inhibition of *svep1* function and erythrocyte formation, using *svep1* and *gata1* morpholinos, did not lead to any DLAV phenotypes (Figure 2C, D), suggesting that shear stress does not modulate DLAV formation at that developmental stage. However, in the absence of tricaine treatment, complete abolition of blood flow using the *tnnt2a* morpholino lead to a DLAV phenotype in *svep1* morphants (Figure 2E, F) (16.4% ± 16.2 versus 3.5 ± 6.2 in *svep1* morphants only). Abolition of blood flow in svep1 homozygous mutants further increases the proportion of fish exhibiting a strong DLAV phenotype (30% compared to 5.3% of WT clutch mate) (Supplementary Figure 2F). These results suggest that while *svep1* loss-of-function produces a cardiac defect that enhances the effect of tricaine on reducing blood flow, *svep1* has an additive effect in modulating blood vessels anastomosis.

Finally, we observed that *svep1* loss of function significantly increases the percentage of short ISVs in *tnn2a* morphants at 48 hpf (14.6% ± 10.5 versus 4.8% ± 7.6 in *tnnt2a* morphants only). This led us to investigate the importance of *svep1* in the regulation of angiogenic sprout identity and behaviour under reduced flow conditions.

### Svep1 is expressed in neurons of the neural tube and its expression is flow dependent

Imaging of the *svep1* reporter line *Tg(svep1: Gal4FF;UAS:GFP)* at 48 hpf showed strong GFP expression in dorsal epithelial cells, above the neural tube, and in individual neurons of the neural tube (Figure 3A, Supplementary Figure 3A, B). Treatment with 1X tricaine from 30 to 48 hpf led to a significant reduction in *svep1* expression throughout the trunk area, particularly in the neural tube at 48 hpf (Figure 3B). In addition, tricaine treatment between 30 and 48 hpf lead to a significant reduction in *svep1* endogenous expression within the neural tube and ventral somite boundary (supplementary Figure 3C), suggesting that blood flow not only sensitises angiogenic sprouts to *svep1* downregulation but also directly affects *svep1* expression.

**Figure 3.**
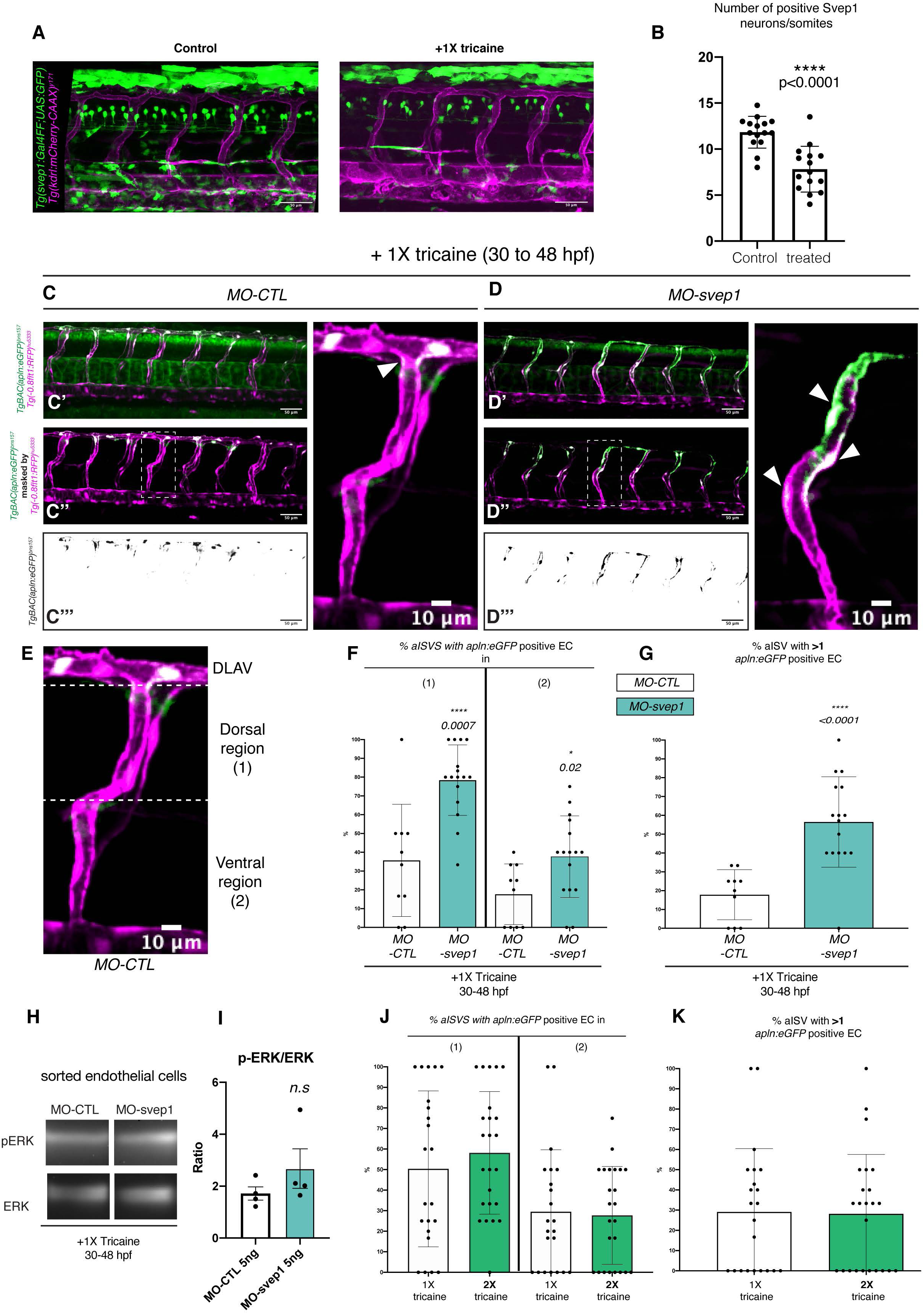
*svep1* loss-of-function leads to a defect in tip/stalk cell specification in primary angiogenic sprouts. **(A)** Representative images of 48 hpf *Tg(svep1:Gal4FF; UAS:eGFP); Tg(kdrl:mcherry-CAAX)^y171^* embryos with or without treatment with 1X (0.014%) tricaine from 30 to 48 hpf. **(B)** Quantification of average numbers of *Tg(svep1:Gal4FF; UAS:eGFP)* positive neurons in the neural tube area of 48 hpf embryos with or without treatment with 1X (0.014%) tricaine from 30 to 48 hpf. (N=3, n=15 controls, n=16 treated). **(C)** Maximum intensity projection of a representative *TgBAC(apln:eGFP)^bns157^, Tg(−0.8flt1:RFP)^hu5333^* MO-CTL (5ng) embryo. C’ shows the unprocessed maximum intensity projection, C’’ shows the GFP signal volume-masked by the RFP signal, to limit detection to the endothelium and C’’’ shows the resulting endothelial GFP signal only. **(D)** Maximum intensity projection of a representative *TgBAC(apln:eGFP)^bns157^, Tg(−0.8flt1:RFP)^hu5333^ MO-CTL* (5ng) embryo. D’ shows the unprocessed maximum intensity projection, D’’ shows the GFP signal volume-masked by the RFP signal, to limit detection to the endothelium and D’’’ shows the resulting endothelial GFP signal only. **(E)** Maximum intensity projection of a MO-CTL (5ng) aISV at 48 hpf, highlighting the ventral and dorsal region used for further quantifications in **(F)** and **(G)**. **(F)** Quantification of the percentage of aISVs with *apln:eGFP* positive endothelial cells in the (1) dorsal and (2) ventral region in 48 hpf MO-CTL (5ng) (n=10) and *MO-svep1*(5ng) (n=16) morphant embryos treated with 1X (0.014%) tricaine from 30 to 48 hpf (N=3). **(G)**Quantification of the percentage of aISVs with more than one *apln:eGFP* positive endothelial cells in 48 hpf *MO-CTL* (5ng) (n =10) and *MO-svep1* (5ng) (n=16) morphant embryos treated with 1X (0.014%) tricaine from 30 to 48 hpf (N=3). **(H)** Representative image of p-ERK and ERK levels in FAC sorted endothelial cells from *MO-CTL* (5ng) and *MO-svep1* (5ng) morphants at 48hpf, treated with 1X (0.014%) tricaine from 30 to 48 hpf (N=4). **(I)** Quantification of p-ERK in FAC sorted endothelial cells of *MO-CTL* (5ng) and *MO-svep1* (5ng) morphants at 48hpf, treated with 1X (0.014%) tricaine from 30 to 48 hpf. Expression levels were normalised to pERK levels (N=4). **(J)** Quantification of the percentage of aISVs with *apln:eGFP* positive endothelial cells in the (1) dorsal and (2) ventral region in 48 hpf embryos treated with 1X (0.014%)(n=22) or 2X (0.028%)(n=24) tricaine from 30 to 48 hpf (N=3). **(K)** Quantification of the percentage of aISVs with more than one *apln:eGFP* positive endothelial cells in 48 hpf embryos treated with 1X (0.014%)(n=22) or 2X (0.028%)(n=24) tricaine from 30 to 48 hpf (N=3).

### *svep1* loss-of-function leads to a defect in tip/stalk cell specification in primary angiogenic sprouts

The formation of the DLAV is initiated by the anastomosis of ipsilateral arterial sprouts, led by a tip cell [1]. To investigate tip cell identity in the zebrafish trunk, we took advantage of the recently published *Tg(apln:eGFP)* [19] reporter line, in which the nucleus of endothelial tip cells is highlighted by eGFP expression at 48 hpf. Following treatment with tricaine from 30 to 48 hpf, *svep1* morphants exhibited an expansion of Apln positive endothelial cells in intersegmental vessels (ISVs), while control morphants predominantly presented Apln positive cells in the dorsal-most region of the vasculature, consistent with a contribution of tip cells to the formation of the DLAV (Figure 3C-F). In addition, *svep1* morphants presented with an overall increase in Apln positive cells per ISVs (56.5% ± 24 versus 17.8% ± 13.3 in control morphants)(Figure 3G).

As tip cell identity is in part characterised by increased levels of p-ERK downstream of Vegfa/Vegfr signalling, we next investigated this pathway activation in *svep1* morphants. Interestingly, sorted endothelial cells from 48 hpf *svep1* morphants treated with 1X tricaine from 30 hpf, did not show a significant increase in p-ERK levels normalised to total ERK levels (Figure 3H, I). Overall, these results suggest that the DLAV phenotype present in *svep1* morphants treated with tricaine is associated with a defect in tip/stalk cell specification in primary angiogenic sprouts.

To test whether the observed expansion of tip cell specification is caused by the augmented reduction in blood flow speed observed in *svep1* loss-of-function embryos, we investigated tip cell identity in control embryos treated with 2X tricaine compared to 1X tricaine. 2X tricaine treatment alone results in an exacerbated blood flow reduction to that observed in *svep1* loss-of-function embryos treated with 1X tricaine (451 um/s ± 231.6, as presented by our laboratory in a recent manuscript [24], versus 521 um/s ± 190). However, treatment with 2X tricaine of Apln-GFP embryos did not cause expansion of tip cell specification (Figure 3J, K), suggesting that this phenotype is primarily a consequence of *svep1* loss-of-function under reduced flow condition.

### *svep1* loss-of-function and knockdown are rescued by *flt1* knockdown

To investigate whether the tip cell phenotype is mediated by, or dependent on, increased Vegfa/Vegfr signaling in *svep1* morphants, we decided to modulate it *in vivo*, first by targeting Flt1 expression. In mice and zebrafish, Flt1 mainly functions as a decoy receptor with high affinity for Vegfa during development to modulate the activation of the Vegfa/Vegfr signalling pathway [21, 25, 26]. Alternative splicing of *flt1* generates two isoforms: a membrane bound form (mFlt1), and a soluble form (sflt1, an alternative spliced and secreted form of mFlt1) [27]. Reports have shown that sFlt1 acts as a negative regulator of tip cell formation in the zebrafish trunk [28].

To reduce *flt1* expression, we used a morpholino targeting both mFlt1 and sFlt1 expression. Similar to previous observations [21], *flt1* morphants do not exhibit any DLAV defects at 48 hpf when treated with tricaine from 30 to 48 hpf. However, in *svep1* mutants and morphants, knockdown of *flt1* expression lead to a rescue of DLAV formation defects (Figure 4A, B). Interestingly, *flt1* knockdown rescued the DLAV segment lumenisation phenotype only in *svep1* morphants but not in mutant embryos (Figure 4C, D), suggesting potential differences in the expressivity of the *flt1* knockdown in *svep1* mutant and morphants.

**Figure 4.**
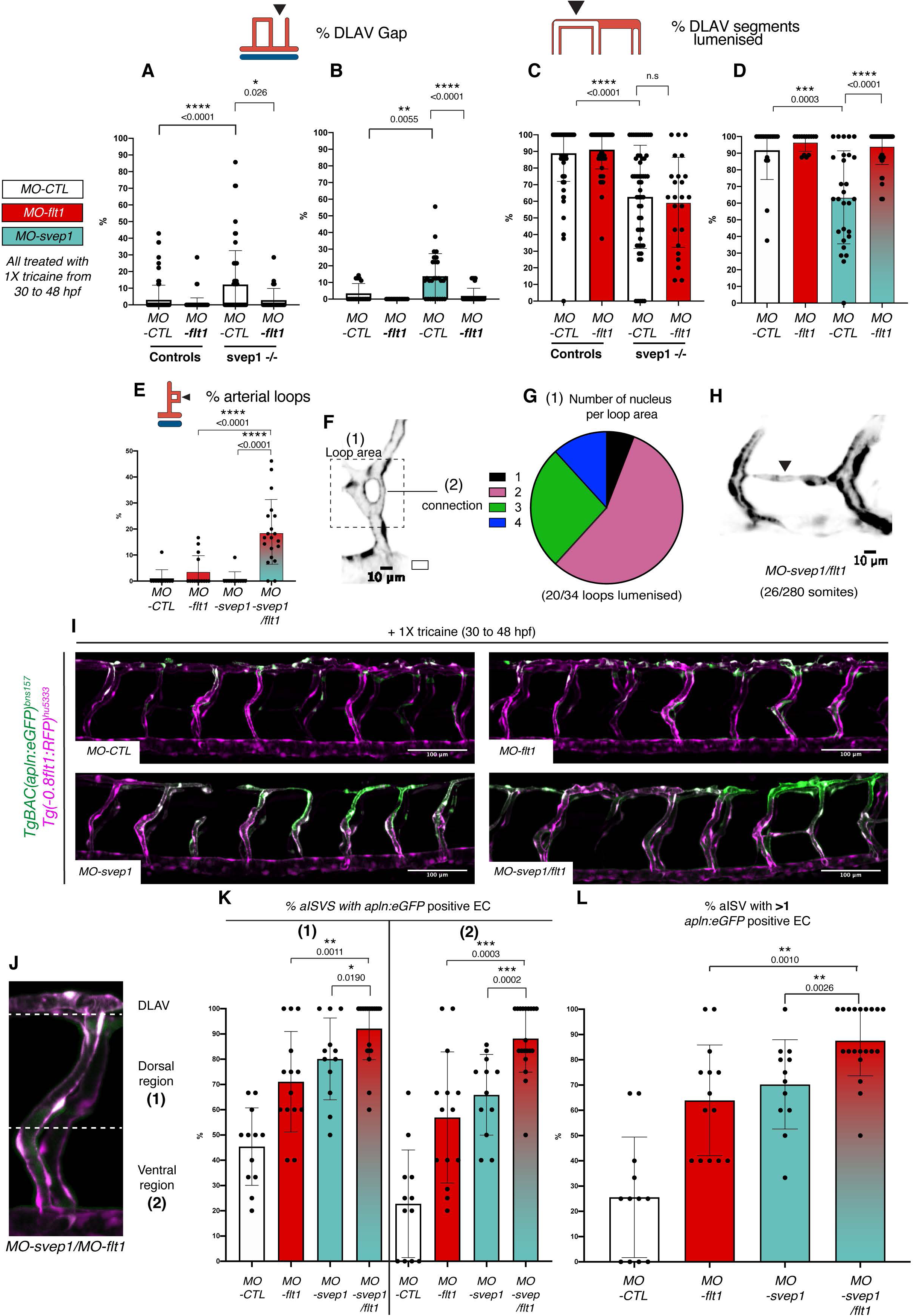
*svep1* loss-of-function and knockdown are rescued by *flt1* knockdown. **(A)** Bilateral quantifications of the percentage of gaps in the DLAV at 48 hpf in controls and mutant *svep1^512-/-^* injected with *MO-CTL* (5ng) (n= 45 and n=27 respectively) or *MO-flt1* (1ng) (n=50 and n=12 respectively), and treated with 1X tricaine (0.014%) from 30 to 48 hpf (N=3). **(B)** Bilateral quantifications of the percentage of gaps in the DLAV at 48 hpf in *MO-CTL* (5 ng) (n=9), *MO-flt1* (1ng) (n=7), *MO-svep1* (5ng) (n=14) and *MO-flt1* (1ng)/*MO-svep1* (5ng) (n=25) embryos (N=3). **(C)** Bilateral quantifications of the percentage of lumenised segments in the DLAV at 48 hpf in controls and mutant *svep1^512-/-^* injected with *MO-CTL* (5ng) (n= 45 and n=27 respectively) or *MO-flt1* (1ng) (n=50 and n=12 respectively),and treated with 1X tricaine (0.014%) from 30 to 48 hpf (N=3). **(D)** Bilateral quantifications of the percentage of lumenised segments in the DLAV at 48 hpf in *MO-CTL* (5 ng) (n=9), *MO-flt1* (1ng) (n=7), *MO-svep1* (5ng) (n=14) and *MO-flt1* (1ng)/MO-svep1 (5ng) (n=25) embryos, treated with 1X tricaine (0.014%) from 30 to 48 hpf (N=3). **(E)** Bilateral quantifications of the percentage of aISV loops at 48 hpf in *MO-CTL* (5 ng) (n=11), *MO-flt1* (1ng) (n=14), *MO-svep1* (5ng) (n=11) and *MO-flt1* (1ng)/*MO-svep1* (5ng) (n=20) embryos treated with 1X tricaine (0.014%) from 30 to 48 hpf (N=3). **(F)** Representative image of an arterial aISV loop in *MO-svep1*(5ng)/*MO-flt1*(1ng) *Tg(−0.8flt1:RFP)^hu3333^* embryos at 48 hpf, treated with 1X(0.014%) tricaine from 30 to 48 hpf. Tg(−0.8flt1:RFP)hu333 **(G)**Quantification of number of nucleus per loop area (see figure 4F) at 48 hpf in *MO-svep1*(5ng)/*MO-flt1*(1ng) embryos at 48 hpf, treated with 1X (0.014%) tricaine from 30 to 48 hpf (n=34 loops counted. 2, 19, 9 and 4 loops had 1, 2, 3 or 4 nucleus per loop area, respectively). 20/34 loops were lumenised) (N=3) **(H)** Representative image of an aISV to aISV connection in the region of the horizontal myoseptum at 48 hpf in in *MO-svep1*(5ng)/*MO-flt1*(1ng) *Tg(−0.8flt1:RFP)^hu3333^* embryos at 48 hpf, treated with 1X (0.014%) tricaine from 30 to 48 hpf. (n=20 fish, 26 connections visible out of 280 somites, 8/26 connections were lumenised, N=3). **(I)** Maximum intensity projection of a representative *TgBAC(apln:eGFP)^bns157^, Tg(−0.8flt1:RFP)^hu5333^* morphant embryos at 48 hpf. The panels show the GFP signal volume-masked by the RFP signal, to limit detection to the endothelium in *MO-CTL* (5ng), *MO-flt1* (1ng), *MO-svep1* (5ng) and *MO-svep1* (5ng) /*MO-flt1* (1ng) embryos treated with 1X (0.014%) tricaine from 30 to 48 hpf. **(J)** Maximum intensity projection of a *MO-svep1* (5ng)/*MO-flt1* (1ng) aISV at 48 hpf, highlighting the ventral and dorsal region used for further quantifications in **(K)**. **(K)** Quantification of the percentage of aISVs with *apln:eGFP* positive endothelial cells in the (1) dorsal and (2) ventral region in 48 hpf *MO-CTL* (5ng) (n=12), *MO-flt1* (1ng) (n=14), *MO-svep1* (5ng)(n=12) and *MO-svep1*(5ng)/*MO-flt1* (1ng) (n=20) morphant embryos treated with 1X (0.014%) tricaine from 30 to 48 hpf (N=3). **(L)** Quantification of the percentage of aISVs with more than one *apln:eGFP* positive endothelial cells in 48 hpf *MO-CTL* (5ng) (n=12),*MO-flt1* (1ng) (n=14), *MO-svep1* (5ng)(n=12) and *MO-svep1* (5ng)/*MO-flt1* (1ng) (n=20) morphant embryos treated with 1X (0.014%) tricaine from 30 to 48 hpf (N=3).

In addition to the DLAV rescue, *flt1/svep1* double knockdown led to the formation of aberrant arterial loops in aISVs (18.5 % ± 12.4 versus 0.8% ± 2.7 in *MO-svep1* and 3.6% ± 6.1 in *MO-flt1* only) (Figure 4E, F). The majority of these arterial loops were lumenised and composed of more than one endothelial cell (>1 cell/loop: 94.1% in n=34 loops). In 9.3% of somites (26/280), we also observed abnormal aISVs to aISVs connections (Figure 4H), which were never seen in control embryos. Thus, the rescue of connectivity at the DLAV level through *flt1* knockdown was accompanied by an excess connectivity at aberrant locations.

Importantly, *flt1* morphants treated with 2X tricaine from 30-48 hpf exhibited aberrant arterial loops in only 2.4% ± 4.1 aISVs, while control morphants under the same treatment never exhibited any (N=3, n=21 *MO-flt1 1ng,* n=24 *MO-CTL 1ng*). In both conditions, we could not find abnormal aISVs to aISVs connections. These results suggest that *svep1* loss-of-function, rather than reduced blood flow, is the principal driver for the excess connectivity observed with concomitant *flt1* knockdown.

In search for the underlying cause of this hyperconnectivity, we investigated tip cell specification using the Apln-GFP transgenic reporter line. In embryos treated with 1X tricaine from 30 to 48hpf, *flt1* knockdown led to an expansion and increase of the total number of Apln+ endothelial cells in aISVs, despite no significant DLAV formation defects (Figure 4I-K). *svep1/flt1* double morphants exhibit significantly more Apln+ endothelial cells in the dorsal and ventral part of aISVs, and more Apln+ cells per aISV than both *svep1* and *flt1* morphants alone (more than one Apln+ cell in 87.6% ± 13.9 ISVs versus 70.3% ± 17.7 in *svep1* only morphants and 63.9 ± 21.9 in *flt1* only morphants)(Figure 1L).

These results suggest that the expansion of the number of tip cells in aISVs per se is not the driver of anastomosis defects.

### Vegfa/Vegfr signalling is necessary for ISV lumenisation maintenance and DLAV formation

Vegfa/Vegfr signalling regulates primary angiogenic sprouting in the developing zebrafish trunk. Inhibition of Vegfr tyrosine kinase activation between 18 and 20 hpf results in the absence of angiogenic sprouting from the dorsal aorta [29]. However, to our knowledge, there exist no reports for the role of active Vegfa/Vegfr signalling in the initial formation and lumenisation of the DLAV (30-48 hpf). As a polarised increase in Vegfa/Vegfr signalling is necessary to establish tip and stalk cells identity in the growing sprouts [6] and a local increase in VEGFA signalling is essential for the establishment of stable connections between vascular sprouts *in vivo* and *in vitro* [30], we decided to investigate the effect of a general reduction of this signalling pathway on the formation of a lumenised DLAV. For this purpose, we used the VEGFR2 inhibitor ZM323881, a tyrosine-kinase inhibitor that reduces Vegfa/Vegfr signalling [31], confirmed by down-regulation of p-ERK in FAC sorted endothelial cells from embryos treated with 50 nM ZM323881 and 1X tricaine from 30 to 48 hpf (Supplementary Figure 3A-B). Consistent with its function, ZM323881-mediated down regulation of Vegfa/Vegfr signalling can be partially rescued with down-regulation of *flt1* expression (Supplementary Figure 3E, F).

Embryos treated with 1X tricaine and 100 or 150 nM ZM323881 from 30 to 48 hpf exhibited significant DLAV defects (Supplementary Figure 3C, D). In addition, even at the highest concentration not inducing any significant defect in DLAV establishment (50nM), we observed a strong defect in ISV lumenisation at 48 hpf (Figure 5D), suggesting that in addition to its importance in the ipsilateral anastomosis of aISVs and DLAV lumenisation, Vegfa/Vegfr signalling is important for the maintenance of aISV lumenisation.

**Figure 5.**
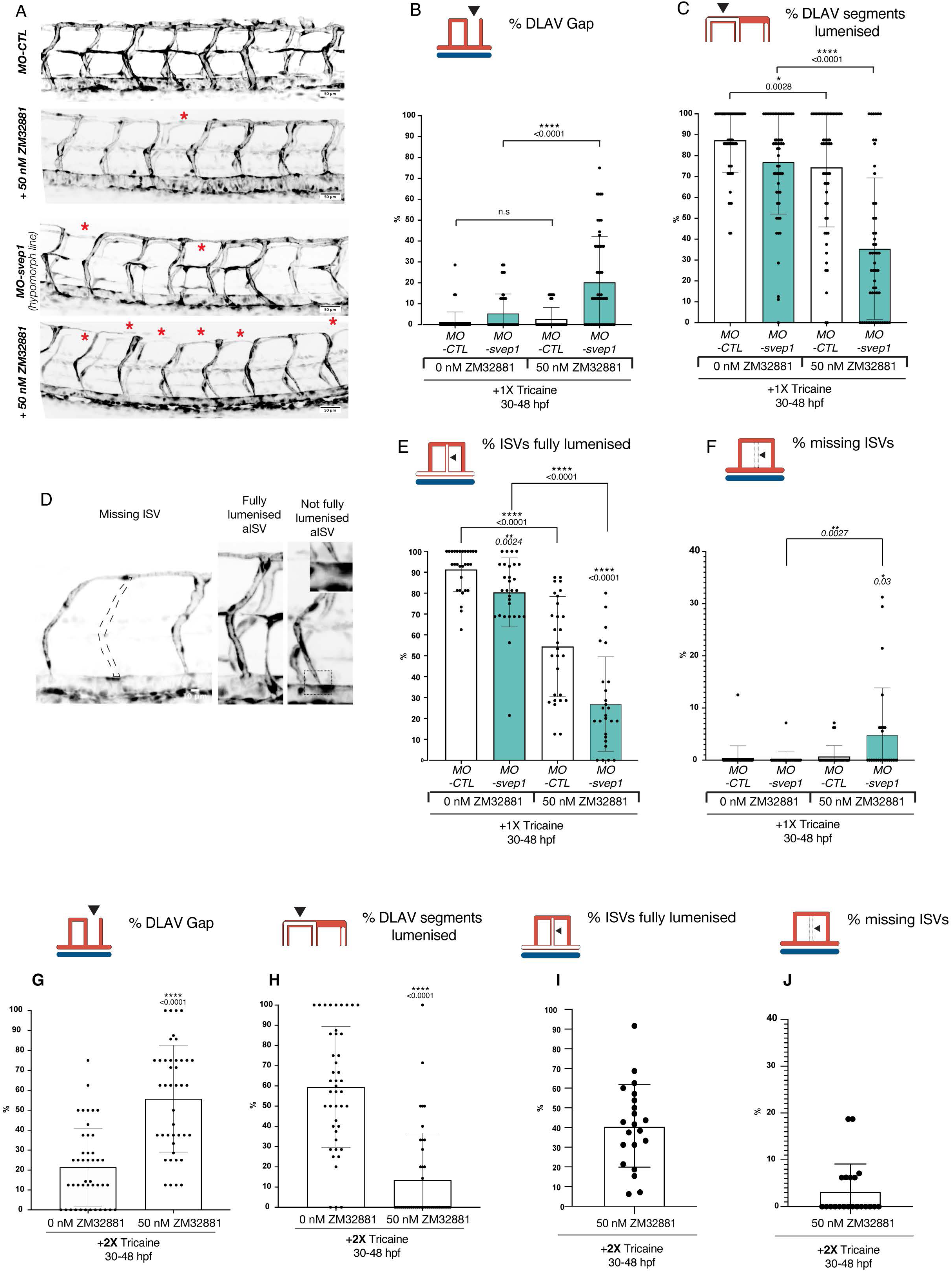
Vegfa/Vegfr signalling is necessary for ISV lumenisation maintenance and DLAV formation. **(A)** Maximum intensity projections at 48 hpf of the trunk of *MO-CTL* (5ng) and *MO-svep1* (5ng), *Tg(fli1a:eGFP)^y7^* embryos, treated with 1X (0.014%) tricaine, with or without 50ng ZM32881. **(B)** Bilateral quantifications of the percentage of gaps in the DLAV at 48 hpf in *MO-CTL* (5 ng) (n=29 (0nM ZM32881), n=28 (50nM ZM32881)) *MO-svep1* (5ng) (n=29 (0 nM ZM32881), n=26 (50nM ZM32881)) embryos treated with 1X (0.014%) tricaine and 0 or 50 nM ZM32881 from 30 to 48 hpf, (N=3). **(C)** Bilateral quantifications of the percentage of lumenised segments in the DLAV at 48 hpf in *MO-CTL* (5 ng) (n=29 (0nM ZM32881), n=28 (50nM ZM32881)) *MO-svep1* (5ng) (n=29 (0 nM ZM32881), n=26 (50nM ZM32881)) embryos treated with 1X (0.014%) tricaine and 0 or 50 nM ZM32881 from 30 to 48 hpf (N=3). **(D)** Representative images of missing ISVs, fully lumenised ISV and not fully lumenised ISV at 48 hpf. Quantifications of these phenotypes are presented in (**E**) and (**F**). **(E)** Bilateral quantifications of the percentage of missing ISVs in the trunk of 48 hpf *MO-CTL* (5 ng) (n=29 (0nM ZM32881), n=28 (50nM ZM32881)) *MO-svep1* (5ng) (n=29 (0 nM ZM32881), n=26 (50nM ZM32881)) embryos treated with 1X (0.014%) tricaine and 0 or 50 nM ZM32881 from 30 to 48 hpf (N=3). **(F)** Bilateral quantifications of the percentage of ISVs lumenised dorsally to ventrally in the trunk of 48 hpf *MO-CTL* (5 ng) (n=29 (0nM ZM32881), n=28 (50nM ZM32881)) *MO-svep1* (5ng) (n=29 (0 nM ZM32881), n=26 (50nM ZM32881)) embryos treated with 1X (0.014%) tricaine and 0 or 50 nM ZM32881 from 30 to 48 hpf (N=3). **(G)**Bilateral quantifications of the percentage of gaps in the DLAV at 48 hpf in embryos treated with 2X (0.028%) tricaine and 0 (n=22) or 50 nM (n=21) ZM32881 from 30 to 48 hpf, (N=3). **(H)** Bilateral quantifications of the percentage of lumenised segments in the DLAV at 48 hpf in embryos treated with 2X (0.028%) tricaine and 0 (n=22) or 50 nM (n=21) ZM32881 from 30 to 48 hpf, (N=3) **(I)** Bilateral quantifications of the percentage of missing ISVs in the trunk of 48 hpf in embryos treated with 2X (0.028%) tricaine and 50 nM (n=21) ZM32881 from 30 to 48 hpf, (N=3). **(J)** Bilateral quantifications of the percentage of ISVs lumenised dorsally to ventrally in the trunk of 48 hpf in embryos treated with 2X (0.028%) tricaine and 50 nM (n=21) ZM32881 from 30 to 48 hpf, (N=3).

### Vegfa/Vegfr signalling inhibition exacerbates *svep1* loss-of-function DLAV phenotype in reduced flow conditions

As the DLAV phenotype in *svep1* morphants can be rescued by an increase in Vegfa/Vegfr signalling (Figure 3), and because an expansion of tip cell numbers within aISVs does not appear to be causative for ipsilateral anastomosis defects (Figure 4), we investigated whether reducing Vegfa/Vegfr signalling in *svep1* morphants would exacerbate the DLAV defect observed under reduced flow conditions.

We took advantage of the variability in expressivity of the DLAV phenotype observed in different reporter lines (Supplementary Figure 1) to select one presenting with a limited DLAV phenotype when injected with *svep1* morpholino and treated with 1X tricaine from 30 to 48 hpf. In this context, we find that concomitant treatment with 50 nM ZM323881 results in a significant increase in the number of gaps in the DLAV (20.3% ± 21.8 versus 5.4% ± 9.3) and in a reduction of the number of lumenised DLAV segments (35.4 ± 33.9 versus 76.9 ± 24.9) (Figure 5A-C). Supporting this notion, we found that treatment with the commonly used VEGFR signalling inhibitor SU5416 [3, 12, 32, 33] also exacerbated the DLAV phenotype in *svep1* morphants treated with 1X tricaine from 30 to 48 hpf (supplementary Figure 5A-C).

In addition, we observed that *svep1* morphants exhibited a mild but significant reduction in ISV lumenisation that was increased by treatment with 50 nM ZM323881 under 1X tricaine conditions (26.9% ± 22.6 versus 80.3% ± 16.5)) (Figure 5E). Furthermore, we found that concomitant *svep1* knockdown and Vegfa/Vegfr signalling inhibition lead to the emergence of embryos with a significant number of ISVs missing at 48 hpf (4.8% ± 9 versus 0.2 ± 1.3%), following treatment with 50 nM ZM323881 and 1X tricaine from 30 to 48 hpf (Figure 5F). Finally, we observed that in the context of Vegfr signalling inhibition, further reducing blood flow with increased tricaine concentration (2X) inhibition did not result in the same phenotypic severity as in *svep1* loss-of-function embryos treated with 1X tricaine. In embryos treated with 50nM ZM323881 and 2X tricaine from 30 to 48 hpf, 40.9% ± 21 of ISVs are lumenised (compared to 26.9% ± 22.6 in mild *svep1 morphants* treated with 1X tricaine and 50nM Zm323881) and 3.3% ± 5.8 of ISVs are missing at 48 hpf (compared to 26.9% ± 22.6 in mild *svep1 morphants* treated with 1X tricaine and 50nM Zm323881). This suggests that reduced function or loss of Svep1 and reduced blood flow both contribute to the emergence of a vascular phenotype in the context of Vegfa/Vegfr signalling inhibition (Figure 5G-J).

Overall, these results suggest that Svep1 and Vegfa/Vegfr signalling appear to act synergistically to maintain vessel lumenisation and stability under low flow conditions.

## Discussion

The question of how vessels anastomose remains incompletely understood, given the complexities of endogenous and exogenous signals driving vascular remodelling and development in parallel and sometimes synergistic ways [1]. Careful and granular analysis of early stage angiogenesis has shed light on the temporal and morphological dynamics underlying this process [11, 34, 35]. Following the establishment of a stable connection between neighbouring tip cells, supported by the local deposition of adherens junctions proteins, such as VE-cadherin, F-actin and ZO-1 [35, 36], ISV connections become progressively lumenised. Lumenisation and stabilisation of ISV connections is thought to occur either through a flow-dependent transcellular hollowing of connecting tip cells (Type I anastomosis)[35, 37] or through a flow-independent process involving the coalescence of isolated luminal pockets into a single luminal space that will subsequently be perfused (Type II anastomosis)[34]. However, comparatively little is known about the molecular pathways leading to the formation and stabilisation of nascent anastomotic connections.

Here, we identified blood flow and Svep1 as regulators of vessel anastomosis in the developing zebrafish vasculature. Both appear to play a role in the stabilisation of nascent anastomotic connections between neighbouring vessels. Under reduced flow conditions, *svep1* knockdown or loss-of-function scenarios result in reduced anastomosis of ipsilateral ISVs and defective formation of a lumenised DLAV at 48 hpf. We find that endothelial cells in *svep1* morphants do not show significantly increased Vegfa/Vegfr signalling. This is surprising given the concomitant increase in Apelin positive cells in ISVs, as tip cells specification has been shown to correlate with higher level of Vegfa/Vegfr signaling [38]. We can speculate that the increase of Vegfa/Vegfr signalling in the trunk ISV of *svep1* morphants is not ubiquitous across the endothelium and therefore cannot be detected with a global endothelial analysis. Alternatively, the increase and expansion of Apln levels in tip cells could be independent of increased ERK activity or Vegfr signalling.

However, neither a Vegfa/Vegr signalling modulation nor the increased tip cell numbers appear causative, as the DLAV defect can be rescued with *flt1* knockdown and is exacerbated by Vegfa/Vegfr signalling inhibition. Interestingly, while further inhibition of blood flow speed below that exhibited in a context of *svep1* loss-of-function and 1X tricaine treatment does similarly result in visible anastomosis defect in the DLAV region, these are not associated with a significant increase of Apelin positive cells in the trunk ISV, supporting our hypothesis that the expansion of the number of tip cells in aISVs per se is not the driver of anastomosis defects.

The results about the role of Vegfa/Vegfr signalling in mediating vessel anastomosis in the zebrafish trunk appear at first sight hard to reconcile. However, previous work on the role of VEGFR2 signalling in anastomosis allow us to speculate on the molecular mechanisms at play here.

In the mouse retina, Nesmith and colleagues have shown that endothelial cells presenting with reduced Flt1 expression were more likely to form stable connections with approaching sprouts [30]. They also clarified that it is reduced mFlt1 expression that influences the bias towards stable connections, suggesting a cell-autonomous regulation of tip cell anastomosis. Finally, they remarked that sprouts with reduced Flt1 expression exhibit reduced exploratory transient connections with adjacent sprouts, proposing that this might lead to an increase in the number of suboptimal new vascular connections. We noticed a significant increase of vascular loops within ISVs, which can functionally be considered suboptimal or redundant for perfusion of tissues, in embryos with reduced *svep1* and *flt1* expression compared to *flt1* knockdown alone. On its own, this result suggests that Svep1 role in vascular development might occur through the modulation of Flt1 activity in vivo. As the function of Flt1 in developmental angiogenesis appears strictly limited to its role as a decoy receptor for VEGFA [26, 39], we can speculate that Svep1 might support/enhance Flt1 decoy ability. Alternatively, we can speculate that *flt1* and *svep1* loss-of-function both drive excess tip cell formation in an additive fashion, and that the DLAV rescue observed in embryos with both loss-of-function could potentially result from an increased tip cell activity, driven by Vegfa/Vegfr signalling increase. This hypothesis would reconcile with our observation that in the context of reduced blood flow, in which embryos present with DLAV formation defects without increase in tip cell numbers, Vegfa/Vegfr signalling increase results in a rescue of the DLAV phenotype without aberrant ISV connections.

However, under reduced flow conditions, while *flt1* knockdown enhances the formation of stable connections between ipsilateral neighbouring sprouts, we find that *svep1 knockdown* alone leads to a significant reduction in the stability of these connections. In addition, we find that inhibition of Vegfa/Vegfr signalling leads to a DLAV phenotype comparable to that observed in *svep1* mutant and morphants, and that Vegfa/Vegfr inhibition in *svep1* morphants leads to a strong increase of anastomosis defects. Taken together, these results suggest that any potential increase in Vegfa/Vegfr signalling following *svep1* knockdown would be compensatory in nature and insufficient to stabilise new connections in the absence of flow.

The importance of blood flow inhibition in the emergence of the DLAV phenotype might suggest a potential investigative avenue. In embryos with reduced blood flow and concomitant reduced Svep1 levels or reduced Vegfa/Vegfr signaling, we observed a significant reduction in ISV lumenisation. This suggest that in this context, a significant proportion of ISVs might initiate anastomosis through flow-independent pathways (Type II anastomosis). We can speculate that blood flow plays a positive role in the stabilization of type II anastomosis between neighboring ISVs, which would explain the DLAV formation defect observed in reduced (tricaine treatment) or abolished (*tnnt2a* morpholino) blood flow conditions. The significant increase in DLAV phenotype observed in *svep1* loss-of-function and abolished flow conditions suggest a role of Svep1 in regulating vessel anastomosis specifically under reduced flow conditions.

In addition, while blood flow reduction between 30 and 48 hpf is sufficient to induce anastomotic defects in the DLAV, this phenotype is significantly exacerbated with concomitant treatment with Vegfr signalling inhibitors, supporting the idea that flow and Vegfa/Vegfr signaling act synergistically to support vessel anastomosis.

Svep1(also known as Polydom) is a secreted ECM protein that has been reported to mediate cell to substrate adhesion in vitro, at leasr in part in an integrin α9β1-dependent manner [13]. Although integrin α9 zebrafish mutants fail to display a vascular phenotype other than lymphatic valve formation [40], interaction with another member of the Integrin family could prove more relevant. For example, Integrin β1b appears to be important for the formation of the DLAV in zebrafish embryos [41]. Future efforts in identifying interaction partners for Svep1 will further enhance our understanding of the molecular pathways regulating vessel anastomosis. Imaging of the *svep1* reporter line *Tg(svep1: Gal4FF;UAS:GFP)* at 48 hpf showed strong GFP expression in dorsal epithelial cells, above the neural tube, and in individual neurons of the neural tube in close proximity to the anastomotic bridges forming between adjacent ISVs. We can speculate that neuronal expression of Svep1 is what locally regulates vessel anastomosis, in a non-cell autonomous manner. In addition, Svep1 is a multi-domain protein, and the importance of individual domains could be functionally tested in reduced flow conditions, by the generation of zebrafish lines expressing selective truncated forms of Svep1. Finally, the significant differences observed in *svep1* loss-of-function phenotype expressivity between zebrafish lines might offer an interesting avenue to decipher the compensatory mechanisms at play and reveal new molecular pathways interacting with Svep1 in the regulation of vessel anastomosis.

Understanding the genetic regulation of phenotypic robustness in angiogenesis and its failure [42] promises crucial insights into the mechanisms causing breakdown of vascular homeostasis in human disease.

## Acknowledgments

We thank all members of the Gerhardt lab for interesting discussions and comments, as well as the Zebrafish facility staff at the MDC for excellent animal care.

## Non-standards Abbreviations and Acronyms

DLAV: dorsal longitudinal anastomotic vessel
hpf: hours post-fertilization
ISVs: intersegmental vessels
DA: Dorsal Aorta
PCV: Posterior Cardinal Vein
VEGFR-2: vascular endothelial growth factor receptor-2
Kdr: Kinase insert domain receptor
Kdrl: Kinase insert domain receptor like
VEGFA: vascular endothelial growth factor A
MO: Morpholino

## References

1. Hogan, B.M. and S. Schulte-Merker, How to Plumb a Pisces: Understanding Vascular Development and Disease Using Zebrafish Embryos. Dev Cell, 2017. 42(6): p. 567–583.

2. Isogai, S., et al., Angiogenic network formation in the developing vertebrate trunk. Development, 2003. 130(21): p. 5281–90.

3. Covassin, L.D., et al., Distinct genetic interactions between multiple Vegf receptors are required for development of different blood vessel types in zebrafish. Proc Natl Acad Sci U S A, 2006. 103(17): p. 6554–9.

4. Covassin, L.D., et al., A genetic screen for vascular mutants in zebrafish reveals dynamic roles for Vegf/Plcg1 signaling during artery development. Dev Biol, 2009. 329(2): p. 212–26.

5. Bussmann, J., et al., Arteries provide essential guidance cues for lymphatic endothelial cells in the zebrafish trunk. Development, 2010. 137(16): p. 2653–7.

6. Shin, M., et al., Vegfa signals through ERK to promote angiogenesis, but not artery differentiation. Development, 2016. 143(20): p. 3796–3805.

7. Bahary, N., et al., Duplicate VegfA genes and orthologues of the KDR receptor tyrosine kinase family mediate vascular development in the zebrafish. Blood, 2007. 110(10): p. 3627–36.

8. Bussmann, J., et al., Zebrafish VEGF receptors: a guideline to nomenclature. PLoS Genet, 2008. 4(5): p. e1000064.

9. Geudens, I. and H. Gerhardt, Coordinating cell behaviour during blood vessel formation. Development, 2011. 138(21): p. 4569–83.

10. Siekmann, A.F. and N.D. Lawson, Notch signalling limits angiogenic cell behaviour in developing zebrafish arteries. Nature, 2007. 445(7129): p. 781–4.

11. Betz, C., et al., Cell behaviors and dynamics during angiogenesis. Development, 2016. 143(13): p. 2249–60.

12. Zygmunt, T., et al., ’In parallel’ interconnectivity of the dorsal longitudinal anastomotic vessels requires both VEGF signaling and circulatory flow. J Cell Sci, 2012. 125(Pt 21): p. 5159–67.

13. Sato-Nishiuchi, R., et al., Polydom/SVEP1 is a ligand for integrin alpha9beta1. J Biol Chem, 2012. 287(30): p. 25615–30.

14. Karpanen, T., et al., An Evolutionarily Conserved Role for Polydom/Svep1 During Lymphatic Vessel Formation. Circ Res, 2017. 120(8): p. 1263–1275.

15. Morooka, N., et al., Polydom Is an Extracellular Matrix Protein Involved in Lymphatic Vessel Remodeling. Circ Res, 2017. 120(8): p. 1276–1288.

16. Kimmel, C.B., et al., Stages of embryonic development of the zebrafish. Developmental dynamics : an official publication of the American Association of Anatomists, 1995. 203(3): p. 253–310.

17. Lawson, N.D. and B.M. Weinstein, In vivo imaging of embryonic vascular development using transgenic zebrafish. Dev Biol, 2002. 248(2): p. 307–18.

18. Traver, D., et al., Transplantation and in vivo imaging of multilineage engraftment in zebrafish bloodless mutants. Nature immunology, 2003. 4(12): p. 1238–46.

19. Marin-Juez, R., et al., Coronary Revascularization During Heart Regeneration Is Regulated by Epicardial and Endocardial Cues and Forms a Scaffold for Cardiomyocyte Repopulation. Dev Cell, 2019. 51(4): p. 503–515 e4.

20. Alestrom, P., et al., Zebrafish: Housing and husbandry recommendations. Lab Anim, 2019: p. 23677219869037.

21. Wild, R., et al., Neuronal sFlt1 and Vegfaa determine venous sprouting and spinal cord vascularization. Nat Commun, 2017. 8: p. 13991.

22. Schindelin, J., et al., Fiji: an open-source platform for biological-image analysis. Nat Methods, 2012. 9(7): p. 676–82.

23. Karpanen, T. and J. Olweus, The Potential of Donor T-Cell Repertoires in Neoantigen-Targeted Cancer Immunotherapy. Front Immunol, 2017. 8: p. 1718.

24. Geudens, I., et al., Artery-vein specification in the zebrafish trunk is pre-patterned by heterogeneous Notch activity and balanced by flow-mediated fine-tuning. Development, 2019. 146(16).

25. Ito, N., et al., Identification of vascular endothelial growth factor receptor-1 tyrosine phosphorylation sites and binding of SH2 domain-containing molecules. J Biol Chem, 1998. 273(36): p. 23410–8.

26. Hiratsuka, S., et al., Flt-1 lacking the tyrosine kinase domain is sufficient for normal development and angiogenesis in mice. Proc Natl Acad Sci U S A, 1998. 95(16): p. 9349–54.

27. Kendall, R.L. and K.A. Thomas, Inhibition of vascular endothelial cell growth factor activity by an endogenously encoded soluble receptor. Proc Natl Acad Sci U S A, 1993. 90(22): p. 10705–9.

28. Krueger, J., et al., Flt1 acts as a negative regulator of tip cell formation and branching morphogenesis in the zebrafish embryo. Development, 2011. 138(10): p. 2111–20.

29. Fish, J.E., et al., Dynamic regulation of VEGF-inducible genes by an ERK/ERG/p300 transcriptional network. Development, 2017. 144(13): p. 2428–2444.

30. Nesmith, J.E., et al., Blood vessel anastomosis is spatially regulated by Flt1 during angiogenesis. Development, 2017. 144(5): p. 889–896.

31. Whittles, C.E., et al., ZM323881, a novel inhibitor of vascular endothelial growth factor-receptor-2 tyrosine kinase activity. Microcirculation, 2002. 9(6): p. 513–22.

32. Fong, T.A., et al., SU5416 is a potent and selective inhibitor of the vascular endothelial growth factor receptor (Flk-1/KDR) that inhibits tyrosine kinase catalysis, tumor vascularization, and growth of multiple tumor types. Cancer Res, 1999. 59(1): p. 99–106.

33. De Angelis, J.E., et al., Tmem2 Regulates Embryonic Vegf Signaling by Controlling Hyaluronic Acid Turnover. Dev Cell, 2017. 40(4): p. 421.

34. Herwig, L., et al., Distinct cellular mechanisms of blood vessel fusion in the zebrafish embryo. Curr Biol, 2011. 21(22): p. 1942–8.

35. Lenard, A., et al., In vivo analysis reveals a highly stereotypic morphogenetic pathway of vascular anastomosis. Dev Cell, 2013. 25(5): p. 492–506.

36. Phng, L.K., F. Stanchi, and H. Gerhardt, Filopodia are dispensable for endothelial tip cell guidance. Development, 2013. 140(19): p. 4031–40.

37. Gebala, V., et al., Blood flow drives lumen formation by inverse membrane blebbing during angiogenesis in vivo. Nat Cell Biol, 2016. 18(4): p. 443–50.

38. Gerhardt, H., et al., VEGF guides angiogenic sprouting utilizing endothelial tip cell filopodia. J Cell Biol, 2003. 161(6): p. 1163–77.

39. Fong, G.H., et al., Role of the Flt-1 receptor tyrosine kinase in regulating the assembly of vascular endothelium. Nature, 1995. 376(6535): p. 66–70.

40. Shin, M., et al., Valves Are a Conserved Feature of the Zebrafish Lymphatic System. Dev Cell, 2019. 51(3): p. 374–386 e5.

41. Iida, A., et al., Integrin beta1 activity is required for cardiovascular formation in zebrafish. Genes Cells, 2018. 23(11): p. 938–951.

42. Kasper, D.M., et al., MicroRNAs Establish Uniform Traits during the Architecture of Vertebrate Embryos. Dev Cell, 2017. 40(6): p. 552–565 e5.

