## Supplementary Figures Legends for "Svep1 stabilizes developmental vascular anastomosis in reduced flow conditions"

**Supplementary figure 1**

**(A)** Bilateral quantifications of the percentage of gaps in the DLAV at 48 hpf in Line A: *Tg(fli1a:eGFP)y1* (N=3, n=18 *MO-CTL* 5ng, n=17 *MO-svep1* 5ng), Line B: *TgBAC(apln:eGFP); Tg(-0.8flt1:RFP)* (N=3, n=10 *MO-CTL* 5ng, n=16 *MO-svep1* 5ng), Line C: *TgBAC(flt1:YFP), Tg(kdrl:mCherry)* (N=3, n=6 *MO-CTL* 5ng, n=17 *MO-svep1* 5ng). The embryos were treated with 1X (0.014%) tricaine from 30 to 48 hpf.

**(B)** Bilateral quantifications of the percentage of lumenised segments in the DLAV at 48 hpf in Line A: *Tg(fli1a:eGFP)y1* (N=3, n=18 *MO-CTL* 5ng, n=17 *MO-svep1* 5ng), Line B: *TgBAC(apln:eGFP); Tg(-0.8flt1:RFP)* (N=3, n=10 *MO-CTL* 5ng, n=16 *MO-svep1* 5ng), Line C: *TgBAC(flt1:YFP), Tg(kdrl:mCherry)* (N=3, n=6 *MO-CTL* 5ng, n=17 *MO-svep1* 5ng). The embryos were treated with 1X (0.014%) tricaine from 30 to 48 hpf.

**(C)** Unilateral quantifications of the percentage of gaps in the DLAV at 48 hpf in *svep1*$^{512}$ homozygous mutants (n=3, 4, 5 embryos) treated with 1X tricaine (0.014%) from 30 to 48 hpf.

**(D)** Unilateral quantifications of the percentage of lumenised segments in the DLAV at 48 hpf in *svep1*$^{512}$ homozygous mutants (n=3, 4, 5 embryos) treated with 1X tricaine (0.014%) from 30 to 48 hpf.

**(E)** Bilateral quantifications of the percentage of gaps in the DLAV at 48 hpf in *svep1*$^{512}$ (n=2, 5, 3 embryos) treated with 1X tricaine (0.014%) from 30 to 48 hpf (N=3).

**(F)** Bilateral quantifications of the percentage of lumenised segments in the DLAV at 48 hpf in *svep1*$^{512}$ homozygous mutants (n=2, 5, 3 embryos) treated with 1X tricaine (0.014%) from 30 to 48 hpf (N=3).

**Supplementary Figure 2**

**(A)** Representative image of a *Tg(-0.8flt1:RFP)*$^{hu333}$, svep1$^{512 -/-}$ embryo treated with 1X (0.014%) tricaine from 30 to 48 hpf and imaged at 48 and 72 hpf.

**(B)** Bilateral quantification of the percentage of gaps at 48 and 72 hpf in the DLAV of svep1$^{512 -/-}$ embryo treated with 1X (0.014%) tricaine from 30 to 48 hpf (n=19, N=3).

**(C)** Quantification of heart rate (heartbeats/minute) in the dorsal aorta of *Tg[gata1a:dsRed]*$^{sd}$ MO-CTL (5ng) or MO-svep1 (5ng) embryos at 48 hpf*,* treated with 1X (0.014% )or 2X *(*0.028% ) tricaine from 30 hpf. (N=4, n=25, 24 MO-CTL (1x, 2X), n=27, 23 MO-svep1 (1X, 2X)).

**(D)** Maximum intensity projection of zebrafish trunk in MO-svep1 (5ng)/Mo-tnnt2a (4ng) embryo at 48 hpf. black arrowheads indicate short ISVs.

**(E)** Bilateral quantification of the percentage of short ISVs in the trunk of 48 hpf MO-CTL (5ng)(n=7), MO-tnnt2a (4ng)(n=12), MO-svep1 (5ng)(n=13) and MO-svep1(5ng)/MO-tnnt2a (4ng)(n=15) embryos (N=3).

**(F)** Percentage of fish presenting with less than 12.5%, between 12.5 and 25% or above 25% gaps in the DLAV at 48 hpf in *svep1*$^{+/+}$ (n=19) , *svep1*$^{+/-}$ (n=35) or *svep* $^{-/-}$ (n=20) embryos injected with *MO-tnnt2a* (4ng) (N=4).

**Supplementary Figure 3**

**(A)** Representative images of 48 hpf *Tg(svep1:Gal4FF; UAS:eGFP); Tg(kdrl:mcherry-CAAX)*$^{y171}$ in the trunk of embryos at 30.5 hpf.

**(B)** Representative images of 48 hpf *Tg(svep1:Gal4FF; UAS:eGFP); Tg(kdrl:mcherry-CAAX)*$^{y171}$ in the trunk of embryos at 48 hpf.

**(C)** *In-situ svep1* endogenous expression in 48hpf embryos not treated (n=11) or treated with 1X (0.014%) tricaine from 30 to 48 hpf (n=18).

**Supplementary Figure 4**

**(A)** Representative image of p-ERK and GFP levels in FAC sorted endothelial cells from embryos treated with 1X (0.014%) tricaine from 30 to 48 hpf and

0nM (n=600 embryos – 290637 cells) or 50 nM (n=600 embryos, 300116 cells) ZM323881(N=3 experiments, pooled).

**(B)** Quantification of p-ERK in FAC sorted endothelial cells from embryos treated with 1X (0.014%) tricaine from 30 to 48 hpf and 0nM (n=600 embryos – 290637 cells) or 50 nM (n=600 embryos, 300116 cells) Expression levels were normalised to GFP levels (N=3 experiments, pooled).

**(C)** Bilateral quantifications of the percentage of gaps in the DLAV at 48 hpf in WT embryos treated with 1X (0.014%) tricaine and 0 (n=16), 50 (n=15), 100 (n=15) and 150 nM (n=15) ZM32881 from 30 to 48 hpf (N=3). Kruskall-Wallis Anova test.

**(D)** Bilateral quantifications of the percentage of lumenised segments in the DLAV at 48 hpf in WT embryos treated with 1X (0.014%) tricaine and 0 (n=16), 50 (n=15), 100 (n=15) and 150 nM (n=15) ZM32881 from 30 to 48 hpf (N=3). Kruskall-Wallis Anova test.

**(E)** Bilateral quantifications of the percentage of gaps in the DLAV at 48 hpf in *MO-CTL* (5 ng) (n=17 (0nM ZM32881), n=16 (150nM ZM32881)) *MO-flt1* (1ng) n=17 (0 nM ZM32881), n=20 (150nM ZM32881)) embryos treated with 1X (0.014%) tricaine and 0 or 50 nM ZM32881 from 30 to 48 hpf (N=4).

**(F)** Bilateral quantifications of the percentage of lumenized segments in the DLAV at 48 hpf in *MO-CTL* (5 ng) (n=17 (0nM ZM32881), n=16 (150nM ZM32881)) *MO-flt1* (1ng) (n=17 (0 nM ZM32881), n=20 (150nM ZM32881)) embryos treated with 1X (0.014%) tricaine and 0 or 50 nM ZM32881 from 30 to 48 hpf (N=4).

**Supplementary Figure 5**

**A)** Representative images of *Tg(fli1a:eGFP)$^{y1}$* embryos at 48 hpf, following treatment with 1X (0.014%) tricaine from 30 hpf, in combination with 0.01% DMSO, 0.1μM SU5416 or 0.25μM SU5416.

**B)** Bilateral quantifications of the percentage of gaps in the DLAV at 48 hpf in *MO-CTL* (5 ng) (n=80 (0.01% DMSO), n=38 (0.1μM SU5416), n=46 (0.25uM DMSO), n=30 (0.5μM SU5416)) *MO-svep1* (5ng) (n=88 (0.01% DMSO), n=44 (0.1μM SU5416), n=30 (0.25μM DMSO), n=22 (0.5μM SU5416)) embryos treated with 1X (0.014%) tricaine and 0.01% DMSO (N=6) or 0.1 (N=3), 0.25 (N=4), 0.5μM (N=4) SU5416 from 30 to 48 hpf.

**C)** Bilateral quantifications of the percentage of lumenised segments in the DLAV at 48 hpf in *MO-CTL* (5 ng) (n=80 (0.01% DMSO), n=38 (0.1μM SU5416), n=46 (0.25uM DMSO), n=30 (0.5μM SU5416)) *MO-svep1* (5ng) (n=88 (0.01% DMSO), n=44 (0.1μM SU5416), n=30 (0.25μM DMSO), n=22 (0.5μM SU5416)) embryos treated with 1X (0.014%) tricaine and 0.01% DMSO (N=6) or 0.1 (N=3), 0.25 (N=4), 0.5μM (N=4) SU5416 from 30 to 48 hpf.

**Supplementary video 1**

Time lapse movie of *MO-CTL* (5ng) *Tg(-0.8flt1:RFP)$^{hu3333}$; TgBAC(flt4:Citrine)* embryo treated with 1X (0.014%) tricaine from 30 to 48 hpf. 15-minute interval between frames.

**Supplementary video 2**

Time lapse movie of *MO-svep1* (5ng) *Tg(-0.8flt1:RFP)$^{hu3333}$; TgBAC(flt4:Citrine)* embryos treated with 1X (0.014%) tricaine from 30 to 48 hpf. 15-minute interval between frames.
