## Supplementary figures and images for "Svep1 stabilizes developmental vascular anastomosis in reduced flow conditions"

### Supplementary Figures

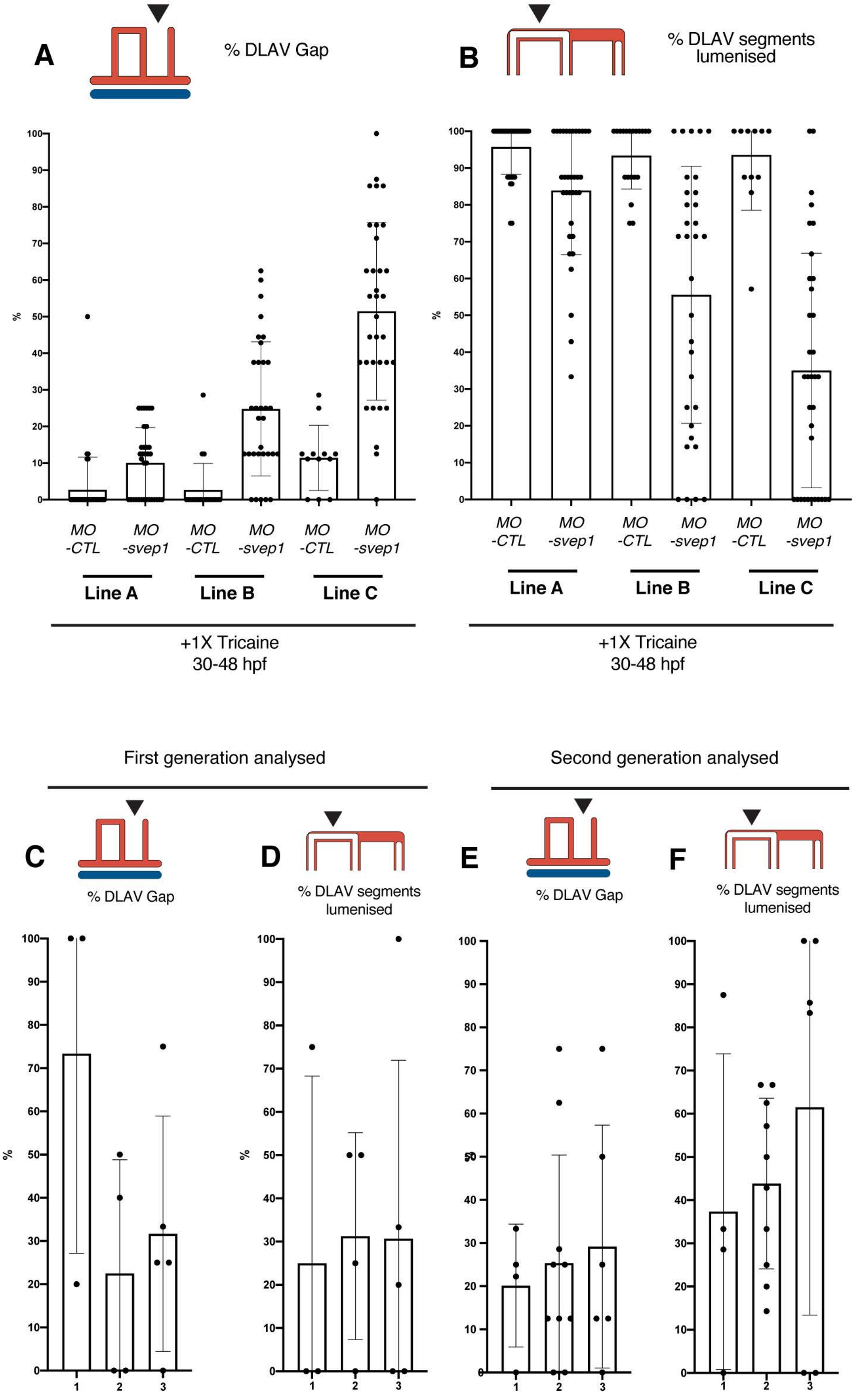

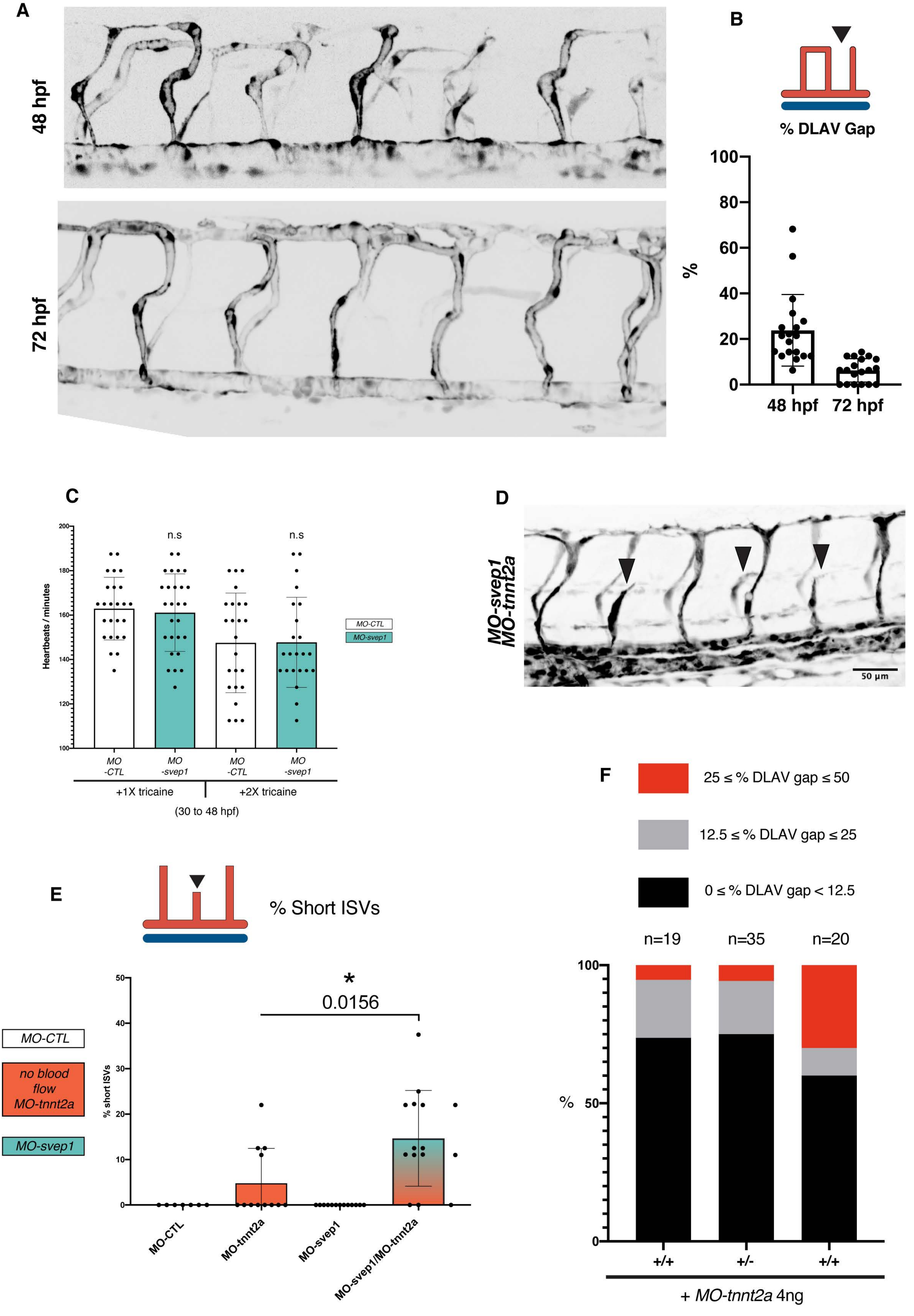

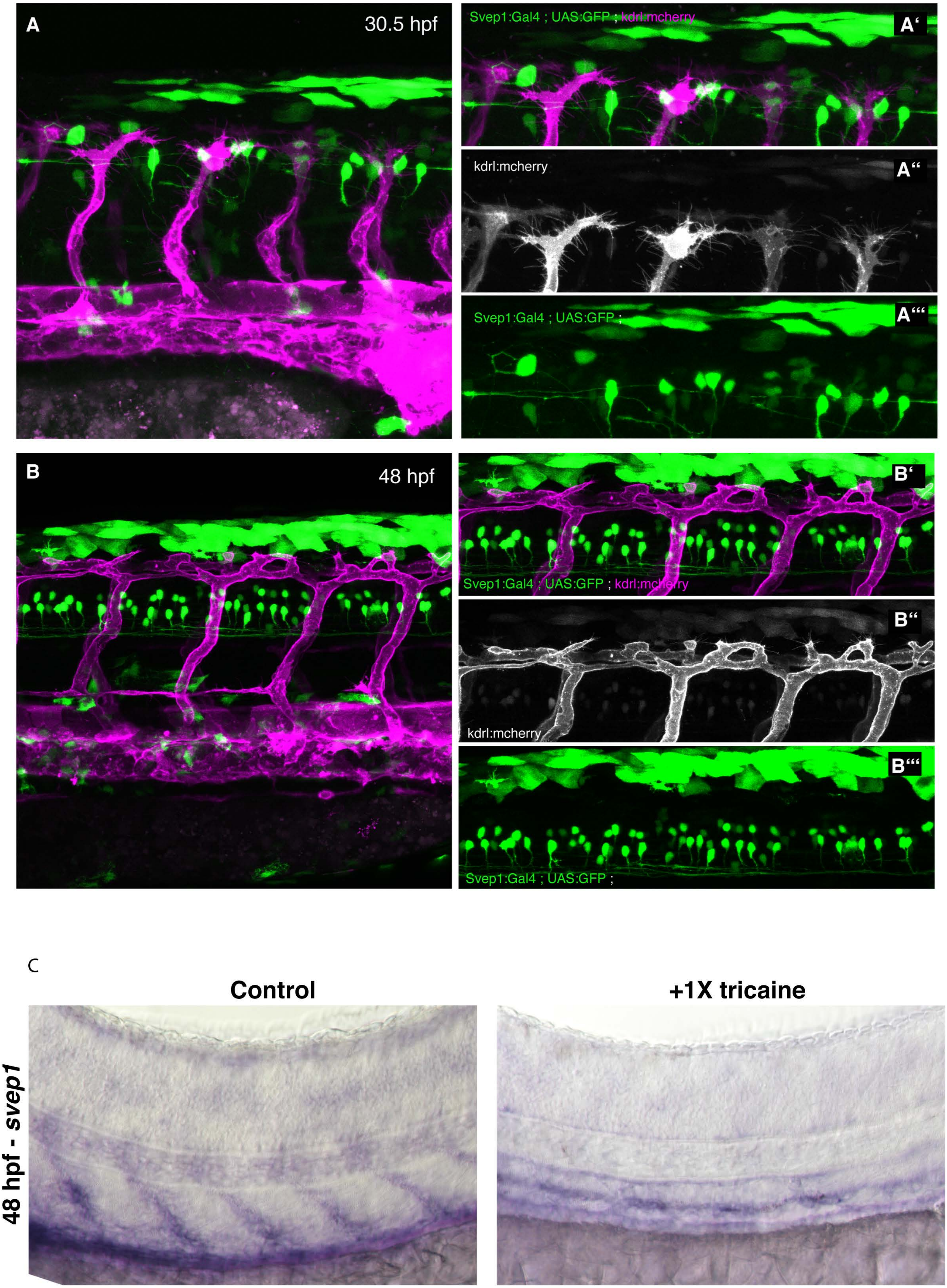

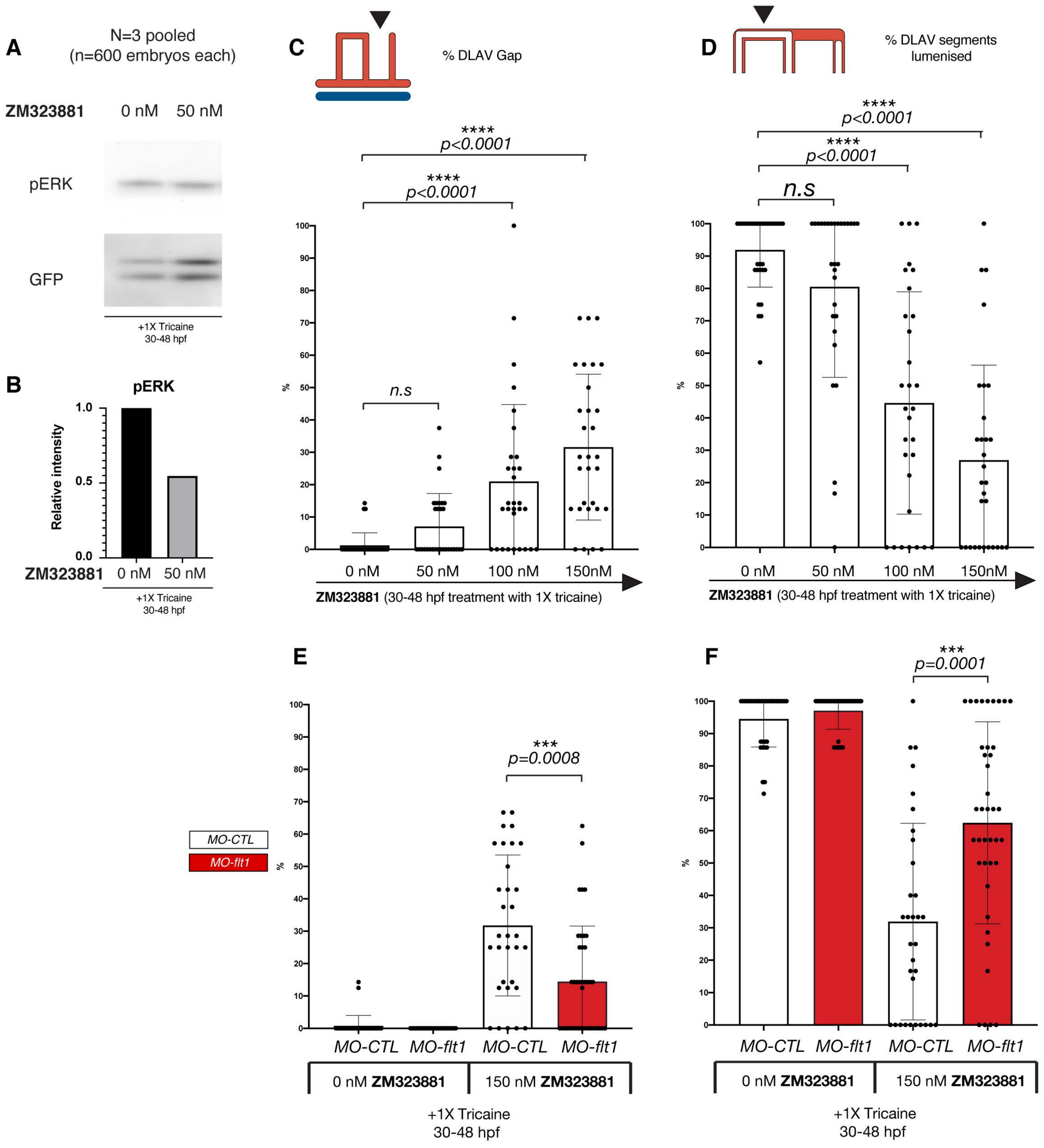

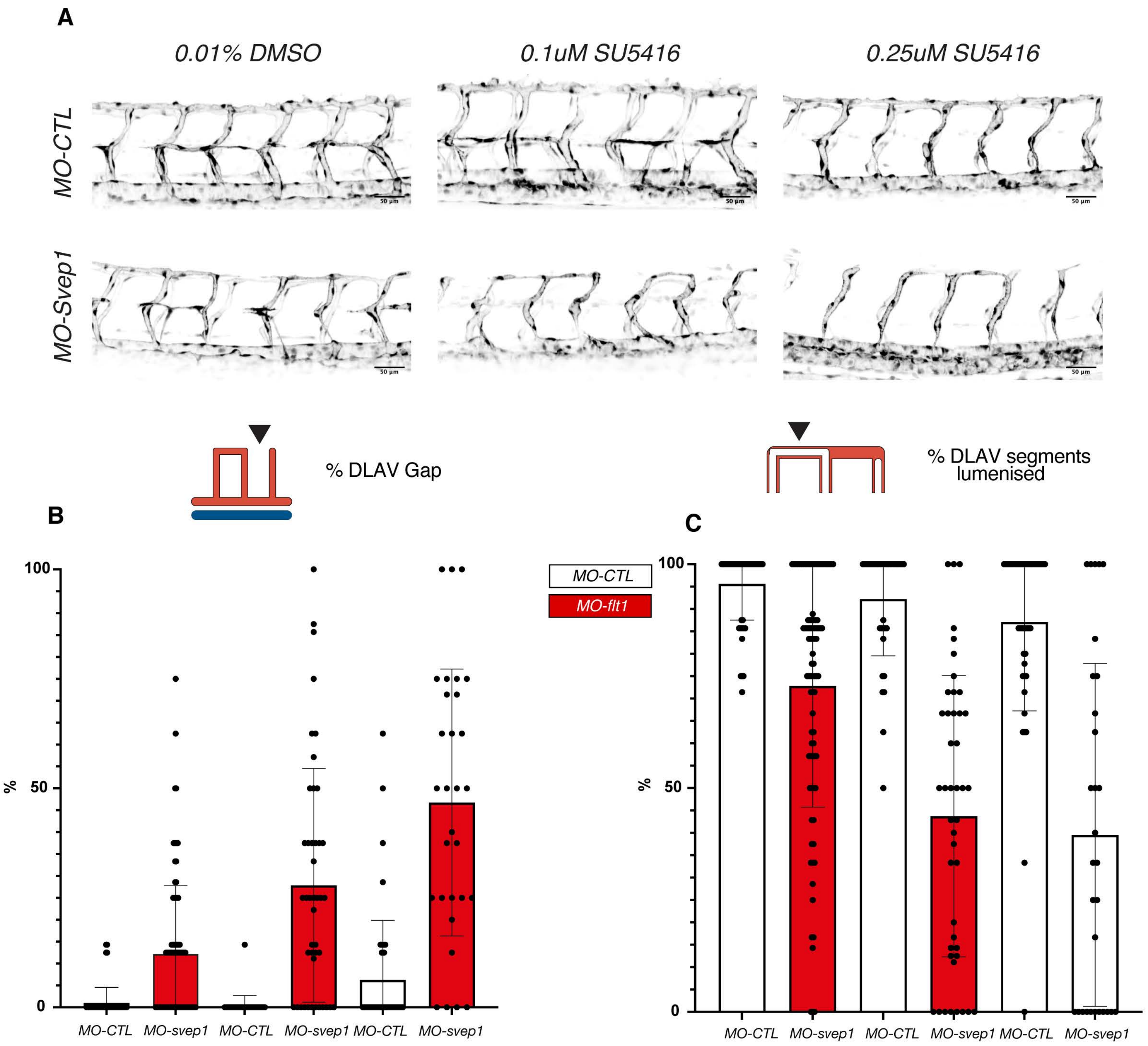
